# AusTraits – a curated plant trait database for the Australian flora

**DOI:** 10.1101/2021.01.04.425314

**Authors:** Daniel Falster, Rachael Gallagher, Elizabeth Wenk, Ian Wright, Dony Indiarto, Caitlan Baxter, Samuel C. Andrew, James Lawson, Stuart Allen, Anne Fuchs, Mark A. Adams, Collin W. Ahrens, Matthew Alfonzetti, Tara Angevin, Owen K. Atkin, Tony Auld, Andrew Baker, Anthony Bean, Chris J. Blackman, Keith Bloomfield, David Bowman, Jason Bragg, Timothy J. Brodribb, Genevieve Buckton, Geoff Burrows, Elizabeth Caldwell, James Camac, Raymond Carpenter, Jane A. Catford, Gregory R. Cawthray, Lucas A. Cernusak, Gregory Chandler, Alex R. Chapman, David Cheal, Alexander W. Cheesman, Si-Chong Chen, Brendan Choat, Brook Clinton, Peta Clode, Helen Coleman, William K. Cornwell, Meredith Cosgrove, Michael Crisp, Erika Cross, Kristine Y. Crous, Saul Cunningham, Ellen Curtis, Matthew I. Daws, Jane L. DeGabriel, Matthew D. Denton, Ning Dong, Honglang Duan, David H. Duncan, Richard P. Duncan, Marco Duretto, John M. Dwyer, Cheryl Edwards, Manuel Esperon-Rodriguez, John R. Evans, Susan E. Everingham, Jennifer Firn, Carlos Roberto Fonseca, Ben J. French, Doug Frood, Jennifer L. Funk, Sonya R. Geange, Oula Ghannoum, Sean M. Gleason, Carl R. Gosper, Emma Gray, Philip K. Groom, Caroline Gross, Greg Guerin, Lydia Guja, Amy K. Hahs, Matthew Tom Harrison, Patrick E. Hayes, Martin Henery, Dieter Hochuli, Jocelyn Howell, Guomin Huang, Lesley Hughes, John Huisman, Jugoslav Ilic, Ashika Jagdish, Daniel Jin, Gregory Jordan, Enrique Jurado, Sabine Kasel, Jürgen Kellermann, Michele Kohout, Robert M. Kooyman, Martyna M. Kotowska, Hao Ran Lai, Etienne Laliberté, Hans Lambers, Byron B. Lamont, Robert Lanfear, Frank van Langevelde, Daniel C. Laughlin, Bree-Anne Laugier-Kitchener, Caroline E. R. Lehmann, Andrea Leigh, Michelle R. Leishman, Tanja Lenz, Brendan Lepschi, James D. Lewis, Felix Lim, Udayangani Liu, Janice Lord, Christopher H. Lusk, Cate Macinnis-Ng, Hannah McPherson, Anthony Manea, Margaret Mayfield, James K. McCarthy, Trevor Meers, Marlien van der Merwe, Daniel Metcalfe, Per Milberg, Karel Mokany, Angela T. Moles, Ben D. Moore, Nicholas Moore, John W. Morgan, William Morris, Annette Muir, Samantha Munroe, Áine Nicholson, Dean Nicolle, Adrienne B. Nicotra, Ülo Niinemets, Tom North, Andrew O’Reilly-Nugent, Odhran S. O’Sullivan, Brad Oberle, Yusuke Onoda, Mark K. J. Ooi, Colin P. Osborne, Grazyna Paczkowska, Burak Pekin, Caio Guilherme Pereira, Catherine Pickering, Melinda Pickup, Laura J. Pollock, Pieter Poot, Jeff R. Powell, Sally A. Power, Iain Colin Prentice, Lynda Prior, Suzanne M. Prober, Jennifer Read, Victoria Reynolds, Anna E. Richards, Ben Richardson, Michael L. Roderick, Julieta A. Rosell, Maurizio Rossetto, Barbara Rye, Paul D. Rymer, Michael A. Sams, Gordon Sanson, Susanne Schmidt, Ernst-Detlef Schulze, Kerrie Sendall, Steve Sinclair, Benjamin Smith, Renee Smith, Fiona Soper, Ben Sparrow, Rachel Standish, Timothy L. Staples, Guy Taseski, Freya Thomas, David T. Tissue, Mark G. Tjoelker, David Yue Phin Tng, Kyle Tomlinson, Neil C. Turner, Erik Veneklaas, Susanna Venn, Peter Vesk, Carolyn Vlasveld, Maria S. Vorontsova, Charles Warren, Lasantha K. Weerasinghe, Mark Westoby, Matthew White, Nicholas Williams, Jarrah Wills, Peter G. Wilson, Colin Yates, Amy E. Zanne, Kasia Ziemińska

## Abstract

We introduce the AusTraits database - a compilation of measurements of plant traits for taxa in the Australian flora (hereafter AusTraits). AusTraits synthesises data on 375 traits across 29230 taxa from field campaigns, published literature, taxonomic monographs, and individual taxa descriptions. Traits vary in scope from physiological measures of performance (e.g. photosynthetic gas exchange, water-use efficiency) to morphological parameters (e.g. leaf area, seed mass, plant height) which link to aspects of ecological variation. AusTraits contains curated and harmonised individual-, species- and genus-level observations coupled to, where available, contextual information on site properties. This data descriptor provides information on version 2.1.0 of AusTraits which contains data for 937243 trait-by-taxa combinations. We envision AusTraits as an ongoing collaborative initiative for easily archiving and sharing trait data to increase our collective understanding of the Australian flora.

## Background and Summary

Species traits are essential metrics for comparing ecological strategies in plants arrayed across environmental space or evolutionary lineages [1, 2, 3, 4]. Broadly, a trait is any measurable property of a plant capturing aspects of its structure or function [5, 6, 7, 8]. Traits thereby provide useful indicators of species’ behaviours in communities and ecosystems, regardless of their taxonomy [8, 9]. Through global initiatives the volume of available trait information for plants has grown rapidly in the last two decades [10, 11]. However, the geographic coverage of trait observations across the globe is patchy, limiting detailed analyses of trait variation and diversity in some regions.

One such region is Australia; a continent with a flora of c. 26,000 native higher-plant species [12]. While significant investment has been made in curating and digitising herbarium collections and observation records in Australia over the last two decades (e.g. The Australian Virtual Herbarium houses ∼7 million specimen occurrence records; https://avh.ala.org.au), no complementary resource yet exists for consolidating information on plant traits. Moreover, relatively few Australian species are represented in the leading global databases. For example, the international TRY database [11] has observations for only 3830 Australian species across all collated traits. This level of species coverage limits our ability to use traits to understand and ultimately manage Australian vegetation [13]. While initiatives such as TRY [11] and the Open Traits Network [14] are working towards global synthesis of trait data, a stronger representation of Australian plant taxa in these efforts is essential given the high richness and endemicity of this continental flora.

Here we introduce the AusTraits database (hereafter AusTraits), a compilation of plant traits for the Australian flora. Currently, AusTraits draws together 351 primary sources and contains 937243 measurements spread across 375 different traits for 29230 taxa. To assemble AusTraits from diverse primary sources and make data available for reuse, we needed to overcome three main types of challenges (Figure 1): 1) Accessing data from diverse original sources, including field studies, online databases, scientific articles, and published taxonomic floras; 2) Harmonising these diverse sources into a federated resource, with common units, trait names, and data formats; and 3) Distributing versions of the data under suitable license. To meet this challenge, we developed a workflow which draws on emerging community standards and our collective experience building trait databases.

**Figure 1:**
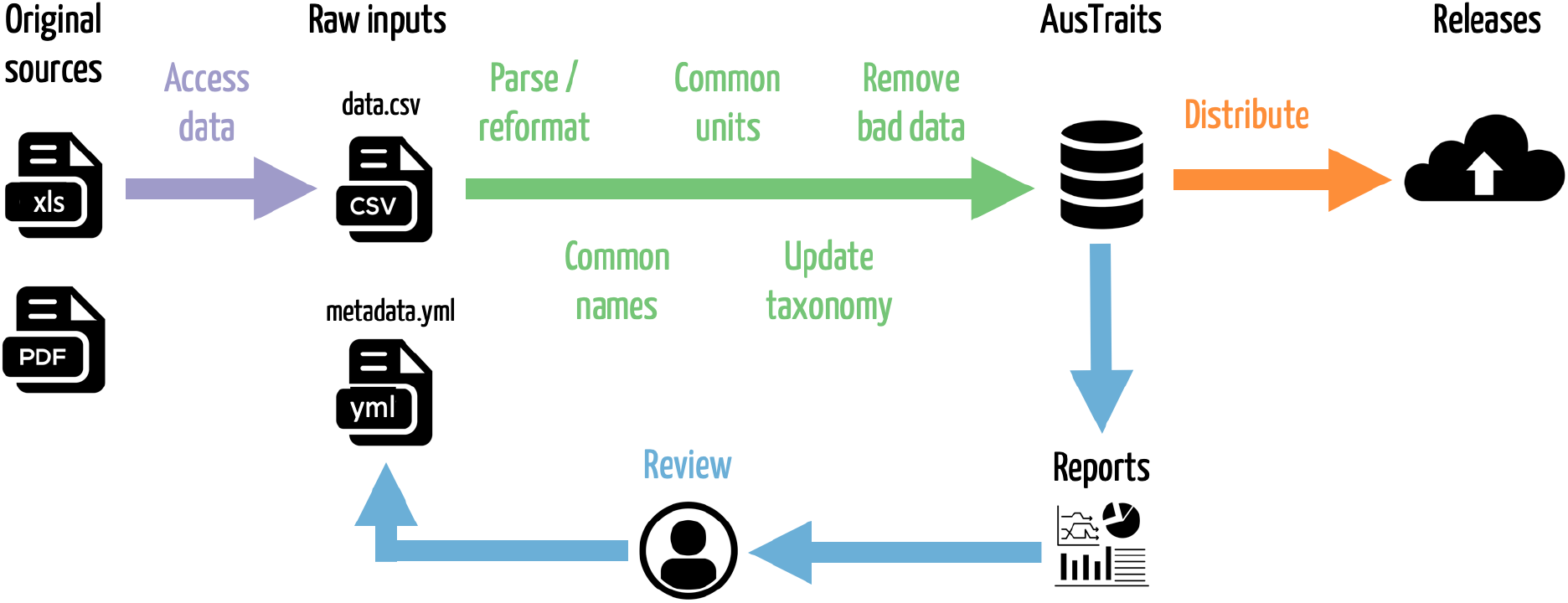
The data curation pathway used to assemble the AusTraits database. Trait observations are accessed from original data sources, including published floras and field campaigns. Features such as variable names, units and taxonomy are harmonised to a common standard. Versioned releases are distributed to users, allowing the dataset to be used and re-used in a reproducible way.

By providing a harmonised and curated dataset on 375 plant traits, AusTraits contributes substantially to filling the gap in Australian and global biodiversity resources. Prior to the development of AusTraits, data on Australian plant traits existed largely as a series of disconnected datasets collected by individual laboratories or initiatives. We envision AusTraits as an on-going collaborative initiative for easily archiving and sharing trait data about the Australian flora. Open access to a comprehensive resource like this will generate significant new knowledge about the Australian flora across multiple scales of interest, as well as reduce duplication of effort in the compilation of plant trait data, particularly for research students and government agencies seeking to access information on traits.

## Methods

### Primary sources

AusTraits version 2.1.0 was assembled from 351 distinct sources, including published papers, field campaigns, botanical collections, and taxonomic treatments (Table 10). Initially we identified a list of candidate traits of interest, then identified primary sources containing measurements for these traits, before contacting authors for access. As the compilation grew, we expanded the list of traits considered to include any measurable quantity that had been quantified for a moderate number of taxa (n > 20).

### Trait definitions

A full list of traits and their sources appears in Table 10 (available online). This list was developed gradually as new datasets were incorporated, drawing from original source publications and a published thesaurus of plant characteristics [15]. We categorised traits based on the tissue where it is measured (bark, leaf, reproductive, root, stem, whole plant) and the type of measurement (allocation, life history, morphology, nutrient, physiological). Version 2.1.0 of AusTraits includes 302 numeric, 71 categorical, and 2 character traits.

### Database schema

The schema of AusTraits broadly follows the principles of the established Observation and Measurement Ontology [16] in that, where available, trait data are connected to contextual information about the collection (e.g. location coordinates, light levels) and information about the methods used to derive measurements (e.g. number of replicates, equipment used). The database contains 11 elements, as described in Table 1. This format was developed to include information about the trait measurements, taxa sampled, the methods used, sites, contextual information, the people involved, and citation sources.

**Table 1:** Main elements of the harmonised AusTraits database. See Tables 2-8 for details on each component.

| Element | Contents |
| --- | --- |
| traits | A table containing measurements of plant traits. |
| sites | A table containing observations of site characteristics associated with information in **traits**. Cross referencing between the two dataframes is possible using combinations of the variables **dataset_id**, **site_name**. |
| contexts | A table containing observations of contextual characteristics associated with information in **traits**. Cross referencing between the two dataframes is possible using combinations of the variables **dataset_id**, **context_name**. |
| methods | A table containing details on methods with which data were collected, including time frame and source. |
| excluded_data | A table of data that did not pass quality test and so were excluded from the master dataset. |
| taxa | A table containing details on taxa associated with information in **traits**. This information has been sourced from the APC (Australian Plant Census) and APNI (Australian Plant Names Index) and is released under a CC-BY3 license. |
| definitions | A copy of the definitions for all tables and terms. Information included here was used to process data and generate any documentation for the study. |
| sources | Bibtex entries for all primary and secondary sources in the compilation. |
| contributors | A table of people contributing to each study. |
| taxonomic_updates | A table of all taxonomic changes implemented in the construction of AusTraits. Changes are determined by comparing against the APC (Australian Plant Census) and APNI (Australian Plant Names Index). |
| build_info | A description of the computing environment used to create this version of the dataset, including version number, git commit and R session_info. |

For storage efficiency, the main table of traits contains relatively little information (Table 2), but can be cross linked against other tables (Tables 3-8) using identifiers for dataset, site, context, observation and taxon (Table 1). The `dataset_id` is ordinarily the surname of the first author and year of publication associated with the source’s primary citation (e.g. `Blackman_2014`). Trait measurements were also recorded as being one of several possible `value_type` (Table 9), reflecting the type of measurement recorded.

**Table 2:** Structure of the `traits` table, containing measurements of plant traits.

| key | value |
| --- | --- |
| dataset_id | Primary identifier for each study contributed into AusTraits; most often these are scientific papers, books, or online resources. By default should be name of first author and year of publication, e.g. **Falster_2005**. |
| taxon_name | Currently accepted name of taxon in the Australian Plant Census or, for unplaced species, in the Australian Plant Names Index. |
| site_name | Name of site where individual was sampled. Cross-references between similar columns in **sites** and **traits**. |
| context_name | Name of contextual senario where individual was sampled. Cross-references between similar columns in **contexts** and **traits**. |
| observation_id | A unique identifier for the observation, useful for joining traits coming from the same **observation_id**. These are assigned automatically, based on the **dataset_id** and row number of the raw data. |
| trait_name | Name of trait sampled. Allowable values specified in the table **traits**. |
| value | Measured value. |
| unit | Units of the sampled trait value after aligning with AusTraits standards. |
| date | Date sample was taken, in the format **yyyy-mm-dd**, but with days and months only when specified. |
| value_type | A categorical variable describing the type of trait value recorded. |
| replicates | Number of replicate measurements that comprise the data points for the trait for each measurement. A numeric value (or range) is ideal and appropriate if the value type is a **mean**, **median**, **min** or **max**. For these value types, if replication is unknown the entry should be **unknown**. If the value type is **raw_value** the replicate value should be 1. If the value type is **expert_mean**, **expert_min**, or **expert_max** the replicate value should be **.na**. |
| original_name | Name given to taxon in the original data supplied by the authors |

**Table 3:** Structure of the `sites` table, containing observations of site characteristics associated with information in `traits`. Cross referencing between the two dataframes is possible using combinations of the variables `dataset_id`, `site_name`.

| key | value |
| --- | --- |
| dataset_id | Primary identifier for each study contributed into AusTraits; most often these are scientific papers, books, or online resources. By default should be name of first author and year of publication, e.g. **Falster_2005**. |
| site_name | Name of site where individual was sampled. Cross-references between similar columns in **sites** and **traits**. |
| site_property | The site characteristic being recorded. Name should include units of measurement, e.g. **longitude (deg)**. Ideally we have at least these variables for each site - **longitude (deg)**, **latitude (deg)**, **description**. |
| value | Measured value. |

**Table 4:**
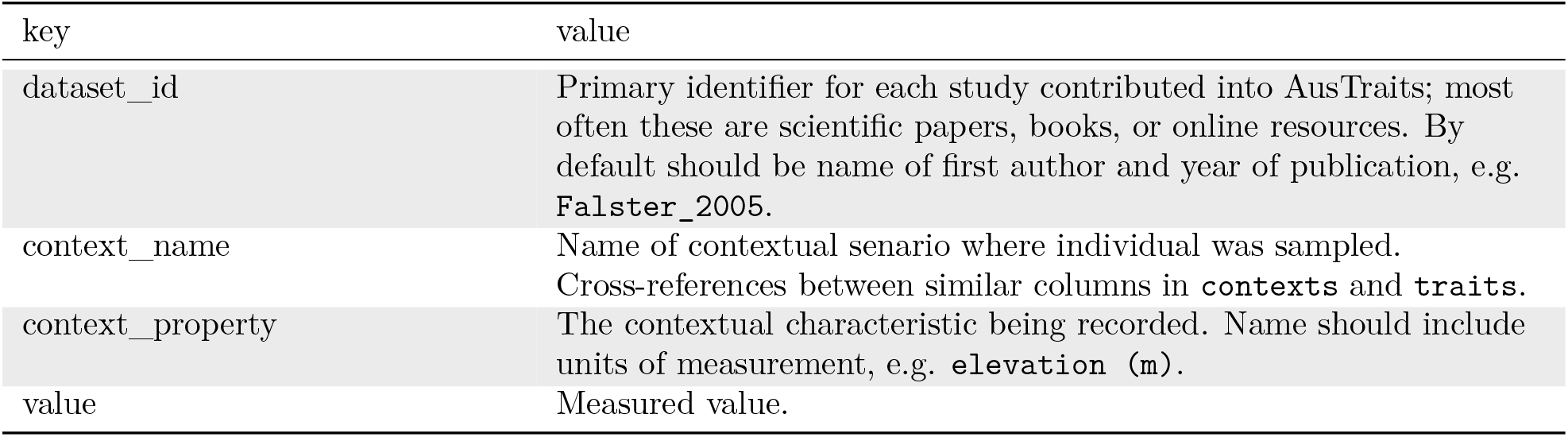
Structure of the `contexts` table, containing observations of contextual characteristics associated with information in `traits`. Cross referencing between the two dataframes is possible using combinations of the variables `dataset_id`, `context_name`.

**Table 5:**
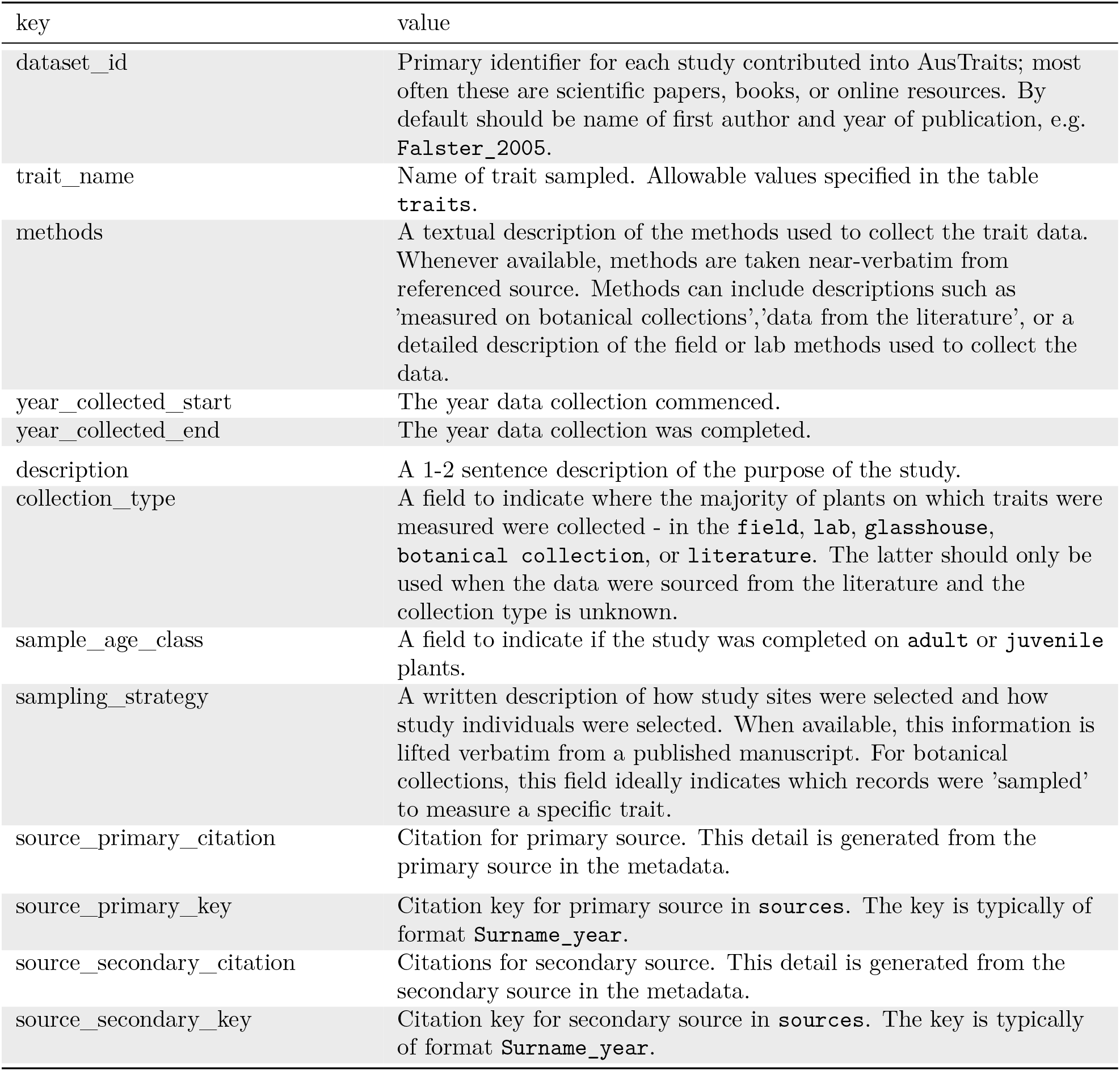
Structure of the `methods` table, containing details on methods with which data were collected, including time frame and source.

| key | value |
| --- | --- |
| dataset_id | Primary identifier for each study contributed into AusTraits; most often these are scientific papers, books, or online resources. By default should be name of first author and year of publication, e.g. **Falster_2005**. |
| trait_name | Name of trait sampled. Allowable values specified in the table **traits**. |
| methods | A textual description of the methods used to collect the trait data. Whenever available, methods are taken near-verbatim from referenced source. Methods can include descriptions such as 'measured on botanical collections', 'data from the literature', or a detailed description of the field or lab methods used to collect the data. |
| year_collected_start | The year data collection commenced. |
| year_collected_end | The year data collection was completed. |
| description | A 1-2 sentence description of the purpose of the study. |
| collection_type | A field to indicate where the majority of plants on which traits were measured were collected - in the **field**, **lab**, **glasshouse**, **botanical collection**, or **literature**. The latter should only be used when the data were sourced from the literature and the collection type is unknown. |
| sample_age_class | A field to indicate if the study was completed on **adult** or **juvenile** plants. |
| sampling_strategy | A written description of how study sites were selected and how study individuals were selected. When available, this information is lifted verbatim from a published manuscript. For botanical collections, this field ideally indicates which records were 'sampled' to measure a specific trait. |
| source_primary_citation | Citation for primary source. This detail is generated from the primary source in the metadata. |
| source_primary_key | Citation key for primary source in **sources**. The key is typically of format **Surname_year**. |
| source_secondary_citation | Citations for secondary source. This detail is generated from the secondary source in the metadata. |
| source_secondary_key | Citation key for secondary source in **sources**. The key is typically of format **Surname_year**. |

**Table 6:** Structure of the `taxonomic_updates` table, of all taxonomic changes implemented in the construction of AusTraits. Changes are determined by comapring against the APC (Australian Plant Census) and APNI (Australian Plant Names Index).

| key | value |
| --- | --- |
| `dataset_id` | Primary identifier for each study contributed into AusTraits; most often these are scientific papers, books, or online resources. By default should be name of first author and year of publication, e.g. **Falster_2005**. |
| `original_name` | Name given to taxon in the original data supplied by the authors |
| `cleaned_name` | Name of the taxon after implementing any changes encoded for this taxon in the metadata file in the specified corresponding **dataset_id**. |
| `taxonIDClean` | Where it could be identified, the **taxonID** of the `cleaned_name` for this taxon in the APC. |
| `taxonomicStatusClean` | Taxonomic status of the taxon identified by **taxonIDClean** in the APC. |
| `alternativeTaxonomicStatusClean` | The status of alternative records with the name `cleaned_name` in the APC. |
| `acceptedNameUsageID` | ID of the accepted name for taxon in the APC or APNI. |
| `taxon_name` | Currently accepted name of taxon in the Australian Plant Census or, for unplaced species, in the Australian Plant Names Index. |

**Table 7:** Structure of the `taxa` table, containing details on taxa associated with information in `traits`. This information has been sourced from the APC (Australian Plant Census) and APNI (Australian Plant Names Index) and is released under a CC-BY3 license.

| key | value |
| --- | --- |
| taxon_name | Currently accepted name of taxon in the Australian Plant Census or, for unplaced species, in the Australian Plant Names Index. |
| source | Source of taxonomic information, either APC or APNI. |
| acceptedNameUsageID | Identifier for the accepted name of the taxon. |
| scientificNameAuthorship | Authority for accepted of the taxon indicated under taxon_name. |
| taxonRank | Rank of the taxon. |
| taxonomicStatus | Taxonomic status of the taxon. |
| family | Family of the taxon. |
| genus | Genus of the taxon. |
| taxonDistribution | Known distribution of the taxon. |
| ccAttributionIRI | Source of taxonomic information. |

**Table 8:** Structure of the `contributors` table, of people contributing to each study.

| key | value |
| --- | --- |
| dataset_id | Primary identifier for each study contributed into AusTraits; most often these are scientific papers, books, or online resources. By default should be name of first author and year of publication, e.g. **Falster_2005**. |
| name | Name of contributor |
| institution | Last known institution or affiliation |
| role | Their role in the study |

**Table 9:** Possible value types of trait records.

| key | value |
| --- | --- |
| raw_value | Value is a direct measurement |
| site_min | Value is the minimum of measurements on multiple individuals of the taxon at a single site |
| site_mean | Value is the mean or median of measurements on multiple individuals of the taxon at a single site |
| site_max | Value is the maximum of measurements on multiple individuals of the taxon at a single site |
| multisite_min | Value is the minimum of measurements on multiple individuals of the taxon across multiple sites |
| multisite_mean | Value is the mean or median of measurements on multiple individuals of the taxon across multiple sites |
| multisite_max | Value is the maximum of measurements on multiple individuals of the taxon across multiple sites |
| expert_min | Value is the minimum observed for a taxon across its range or in this particular dataset, as estimated by an expert based on their knowledge of the taxon. Data fitting this category include estimates from flora that represent a taxon's entire range, and values for categorical variables obtained from a reference book, or identified by an expert. |
| expert_mean | Value is the mean observed for a taxon across its range or in this particular dataset, as estimated by an expert based on their knowledge of the taxon. Data fitting this category include estimates from flora that represent a taxon's entire range, and values for categorical variables obtained from a reference book, or identified by an expert. |
| expert_max | Value is the maximum observed for a taxon across its range or in this particular dataset, as estimated by an expert based on their knowledge of the taxon. Data fitting this category include estimates from flora that represent a taxon's entire range, and values for categorical variables obtained from a reference book, or identified by an expert. |
| experiment_min | Value is the minimum of measurements from an experimental study either in the field or a glasshouse |
| experiment_mean | Value is the mean or median of measurements from an experimental study either in the field or a glasshouse |
| experiment_max | Value is the maximum of measurements from an experimental study either in the field or a glasshouse |
| individual_mean | Value is a mean of replicate measurements on an individual (usually for experimental ecophysiology studies) |
| individual_max | Value is a maximum of replicate measurements on an individual (usually for experimental ecophysiology studies) |
| literature_source | Value is a site or multi-site mean that has been sourced from an unknown literature source |
| unknown | Value type is not currently known |

### Harmonisation

To harmonise each source into the common AusTraits format we applied a reproducible and transparent workflow (Figure 1), written in R [17], using custom code, and the packages `tidyverse` [18], `stringr` [19], `yaml` [20], `remake` [21], `knitr` [22], and `rmarkdown` [23]. In this workflow, we performed a series of operations, including reformatting data into a standardised format, generating observation ids for each individual measured, transforming variable names into common terms, transforming data common units, standardising terms for categorical variables, encoding suitable metadata, and flagging data that did not pass quality checks. Successive versions of AusTraits iterate through the steps in Figure 1, to incorporate new data and correct identified errors, leading to a high-quality, harmonised dataset.

Details from each primary source were saved with minimal modification into two plain text files. The first file, `data.csv`, contains the actual trait data in comma-separated values format. The second file, `metadata.yml`, contains relevant metadata for the study, as well as options for mapping trait names and units onto standard types, and any substitutions applied to the data in processing. These two files provide all the information needed to compile each study into a standardised AusTraits format.

### Taxonomy

We developed a custom workflow to clean and standardise taxonomic names using the latest and most comprehensive taxonomic resources for the Australian flora: the Australian Plant Census (APC) [12] and the Australian Plant Names Index (APNI) [24]. While several automated tools exist, such as `taxize` [25], these do not currently include up to date information for Australian taxa. Updates were completed in two steps. In the first step, we used both direct and then fuzzy matching (with up to 2 characters difference) to search for an alignment between reported names and those in three name sets: 1) All accepted taxa in the APC, 2) All known names in the APC, 3) All names in the APNI. Names were aligned without name authorities, as we found this information was rarely reported in the raw datasets provided to us. Second, we used the aligned name to update any outdated names to their current accepted name, using the information provided in the APC. If a name was recorded as being both an accepted name and an alternative (e.g. synonym) we preferred the accepted name, but also noted the alternative records. When a suitable match could not be found, we manually reviewed near matches and web portals such as the Atlas of Living Australia to find a suitable match. The final resource reports both the original and the updated taxon name alongside each trait record (Table 2), as well an additional table summarising all taxonomic names changes (Table 6) and further information from the APC and APNI on all taxa included (Table 7).

### Data records

#### Access

As an evolving data product, successive versions of AusTraits are being released, containing updates and corrections. Versions are labeled using semantic versioning to indicate the change between versions [26]. Static versions of the AusTraits, including version 2.1.0 used in this descriptor, are available on the project website (http://traitecoevo.github.io/austraits.build/) and Zenodo [27]. The latest data can also be downloaded directly from the project website. As validation (see Technical Validation, below) and data entry is ongoing, users are recommended to pull data from the static releases, to ensure results in their downstream analyses remain consistent as the database is updated.

Data is released under a CC-BY license enabling reuse with attribution – being a citation of this descriptor and, where possible, original sources.

#### Data coverage

The number of accepted vascular plant species in the APC (as of May 2020) is around 24,750 [12]. Version 2.1.0 of AusTraits includes at least one record for 24,148, or about 97% of taxa. Five traits (leaf_length, leaf_width, plant_height, life_history, plant_growth_form) have records for more than 50% of taxa. Across all traits, the median number of taxa with records is 62. Table 10 shows the number of studies, taxa, and families recording data in AusTraits, as well as the number of geo-referenced records, for each trait.

**Table 10:**
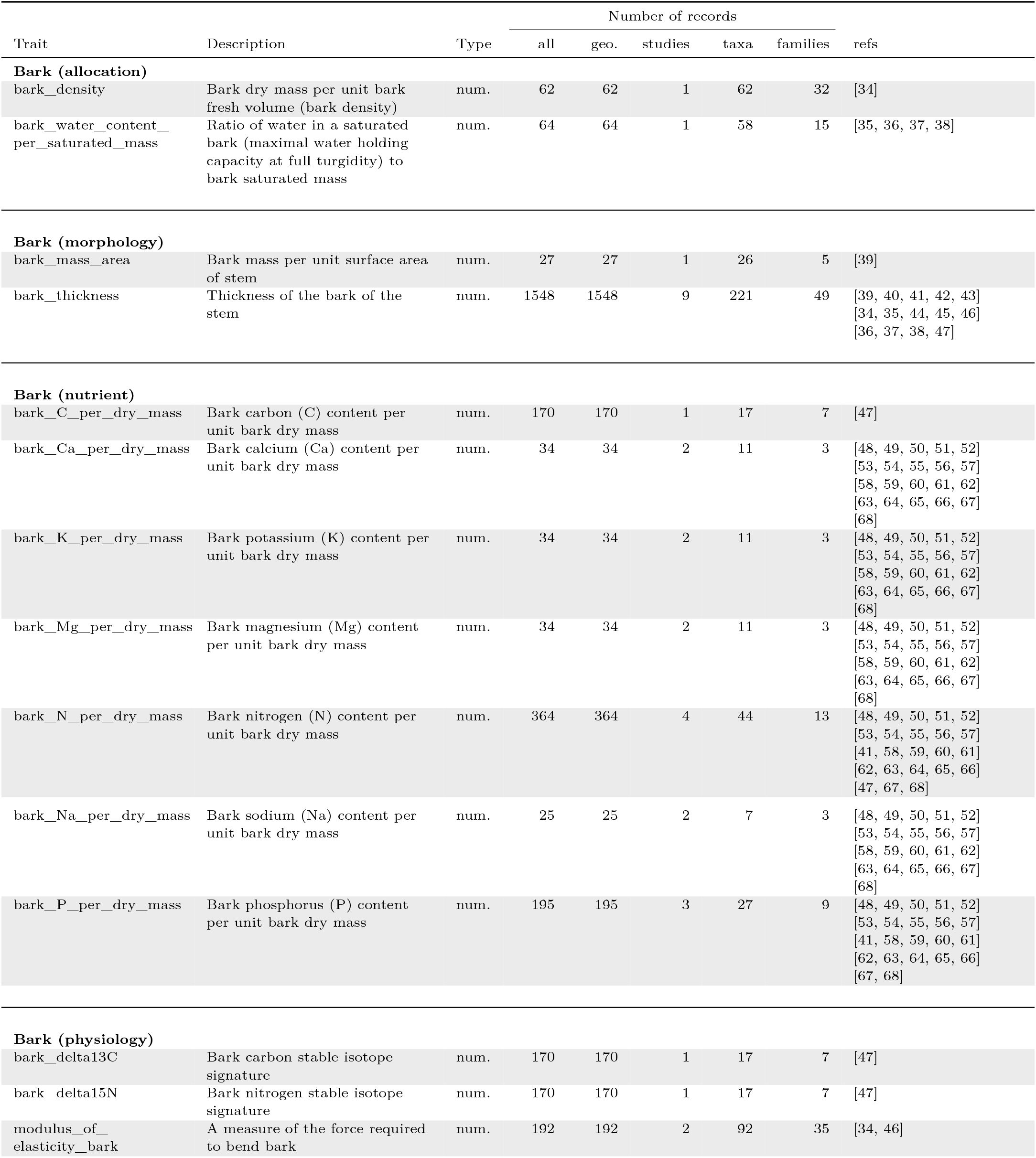
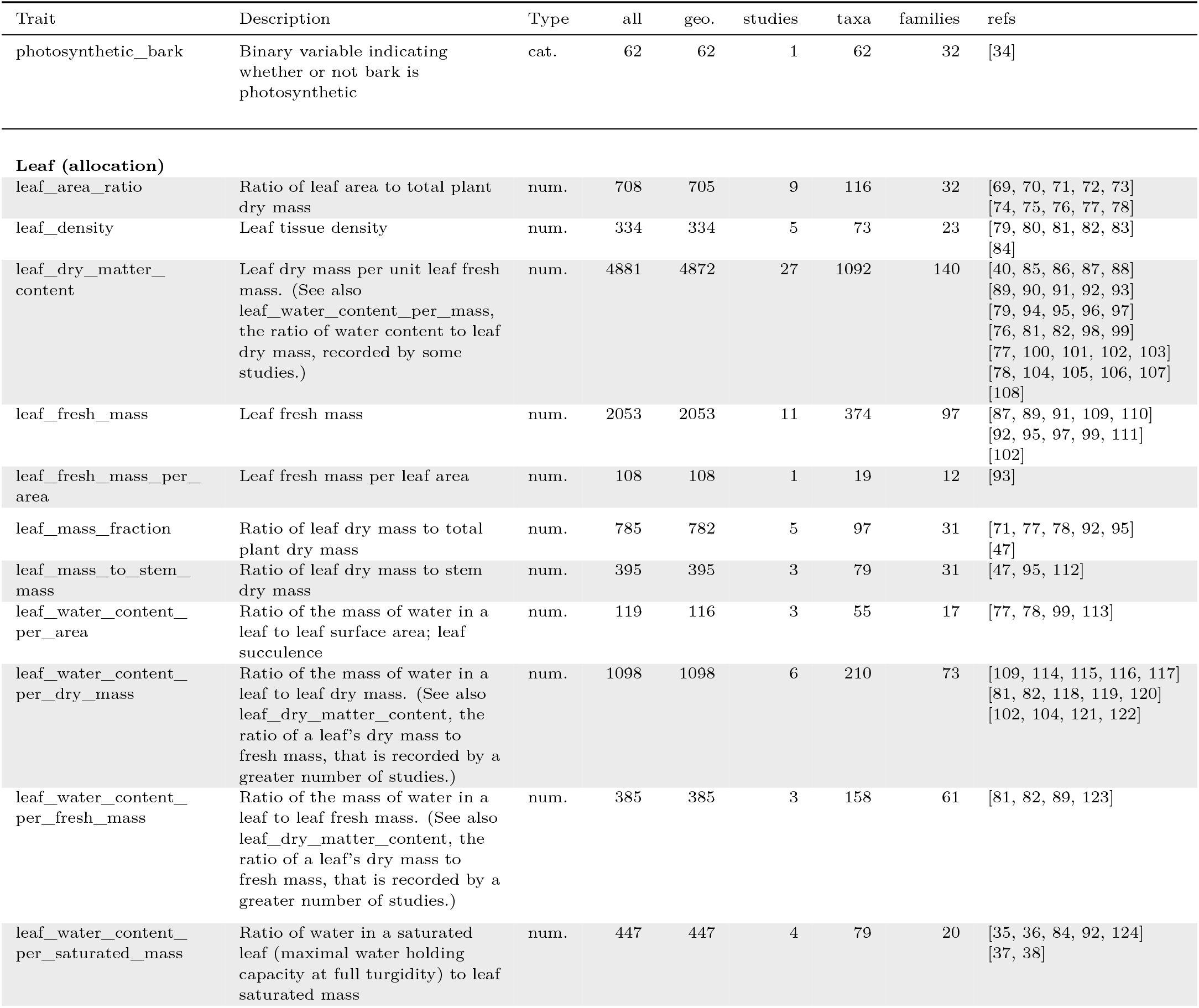
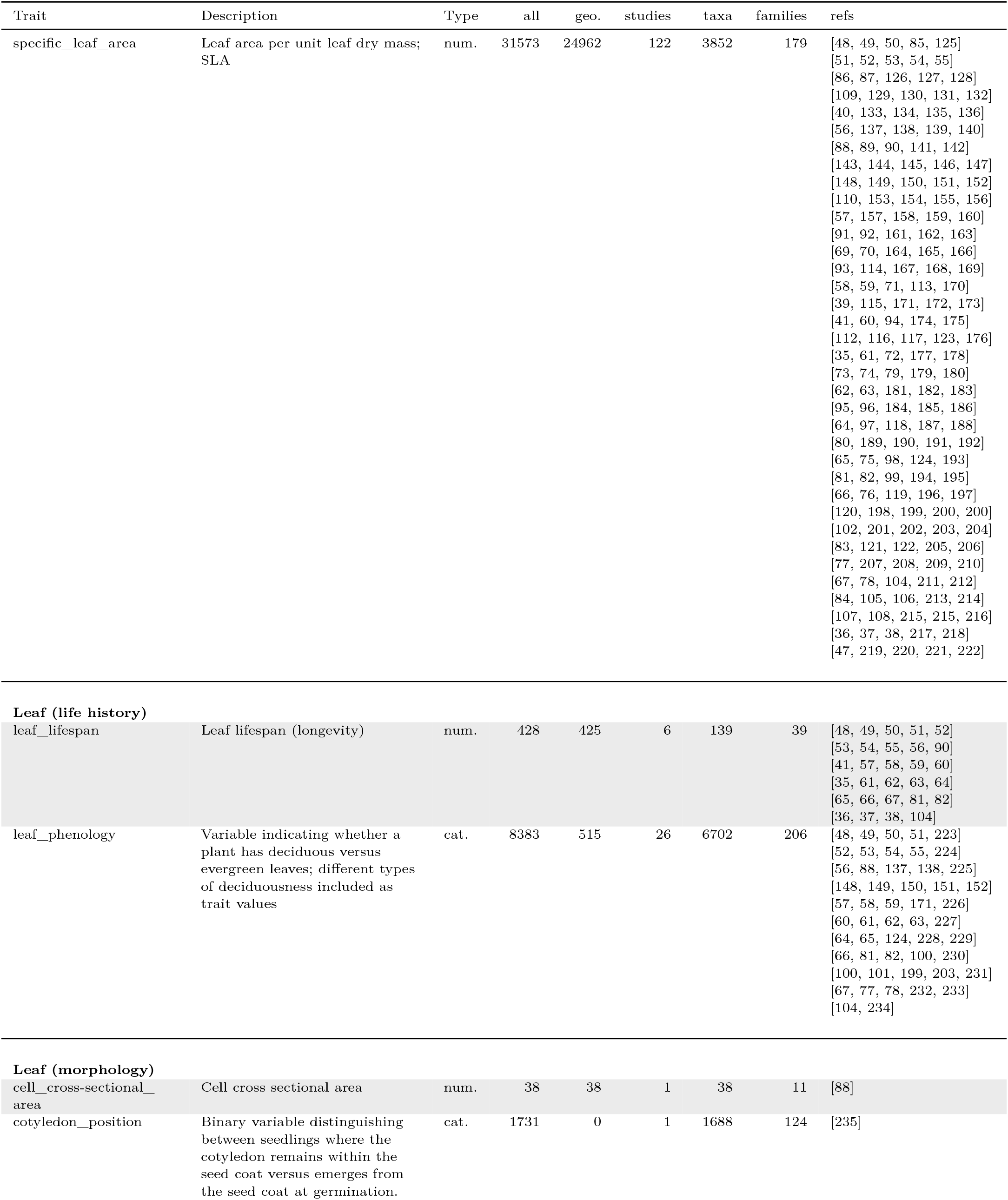
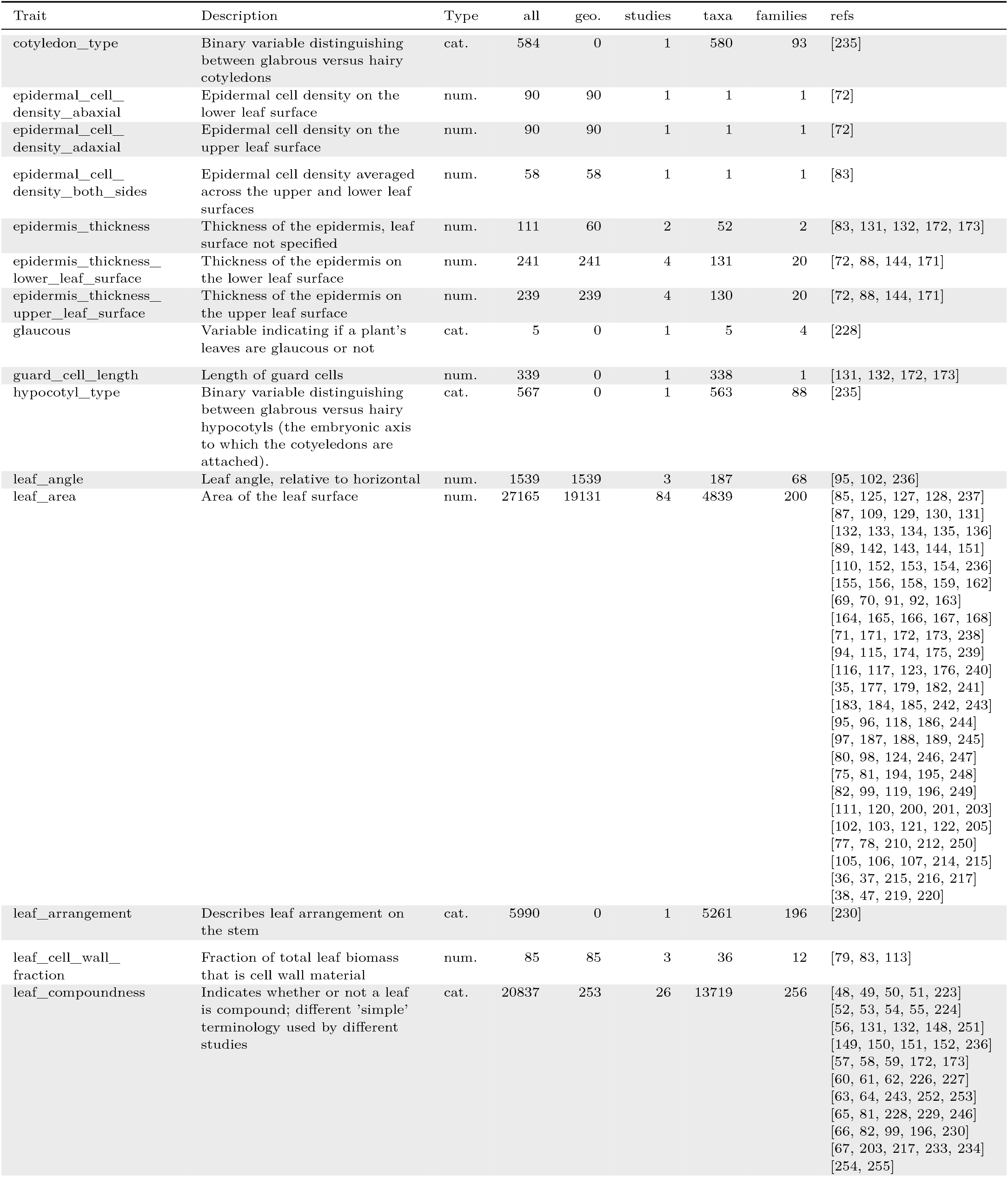
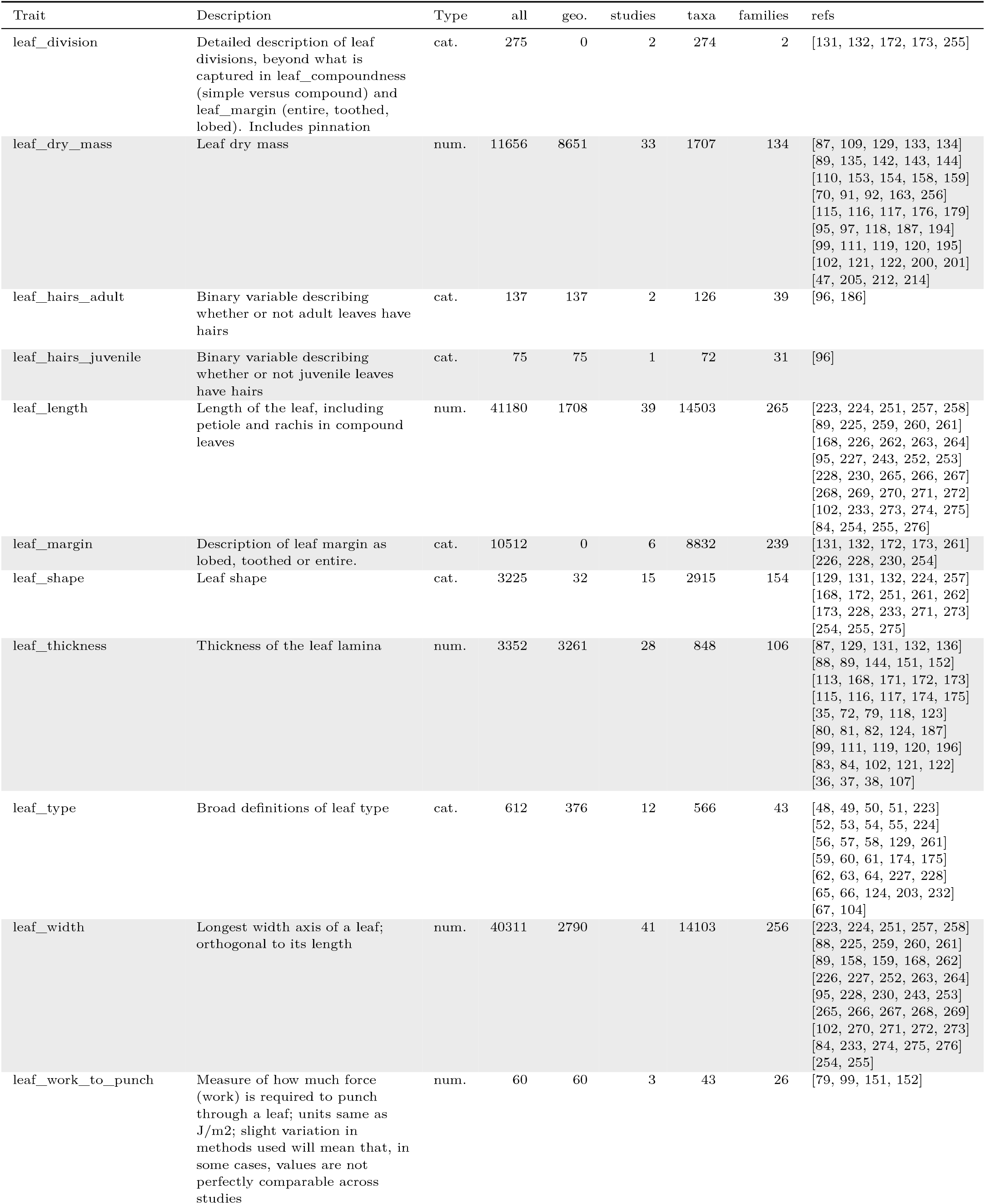
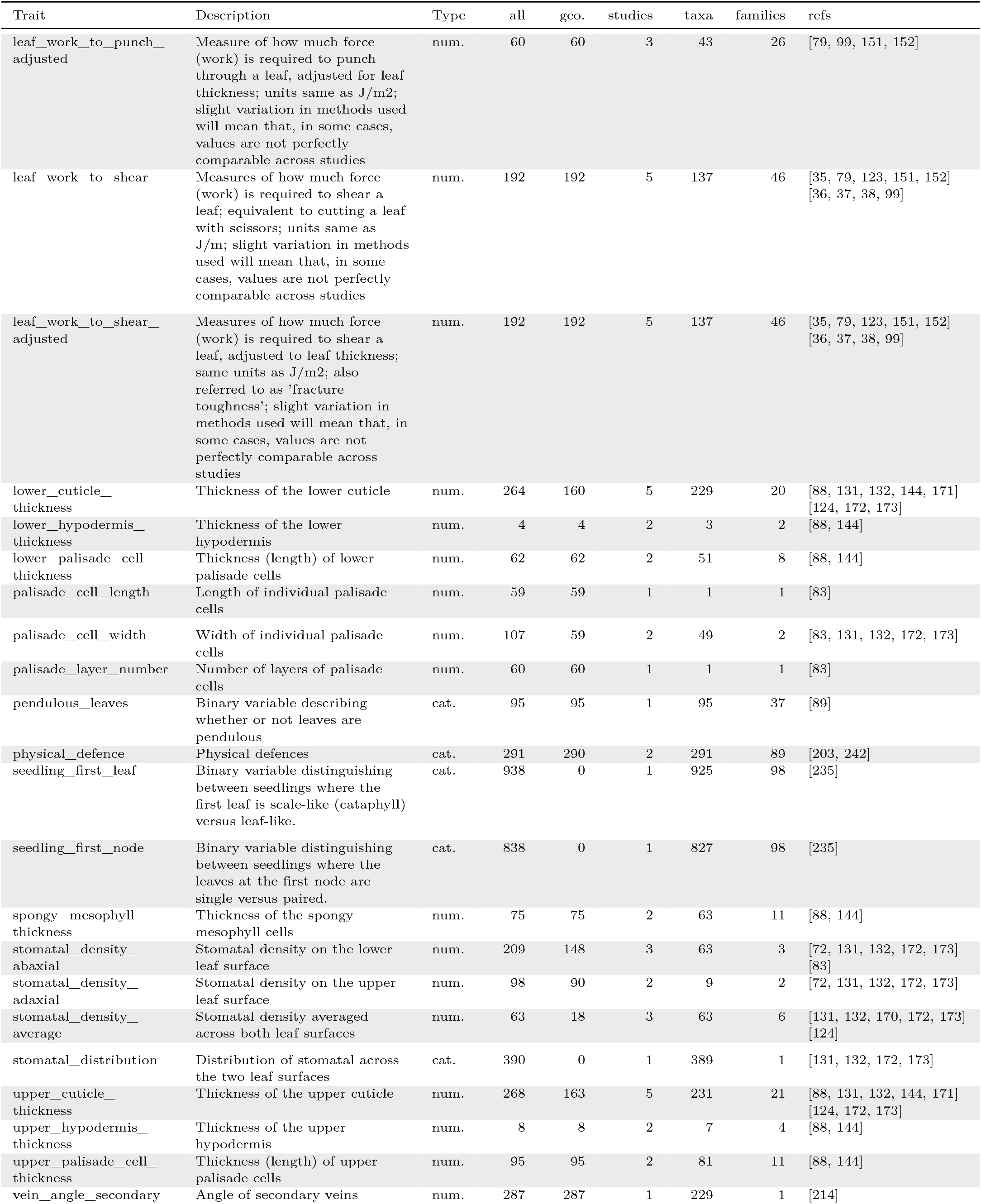
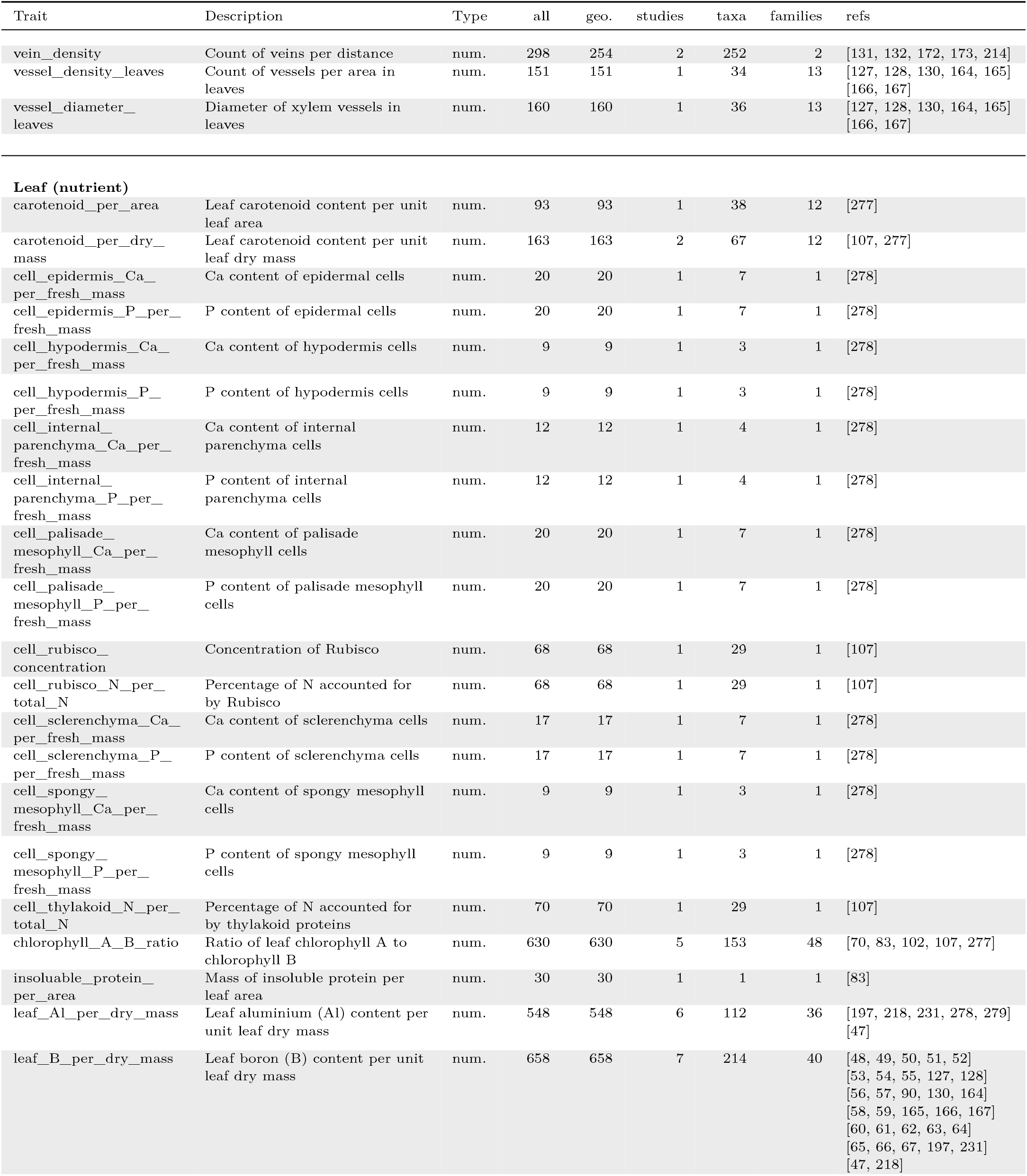
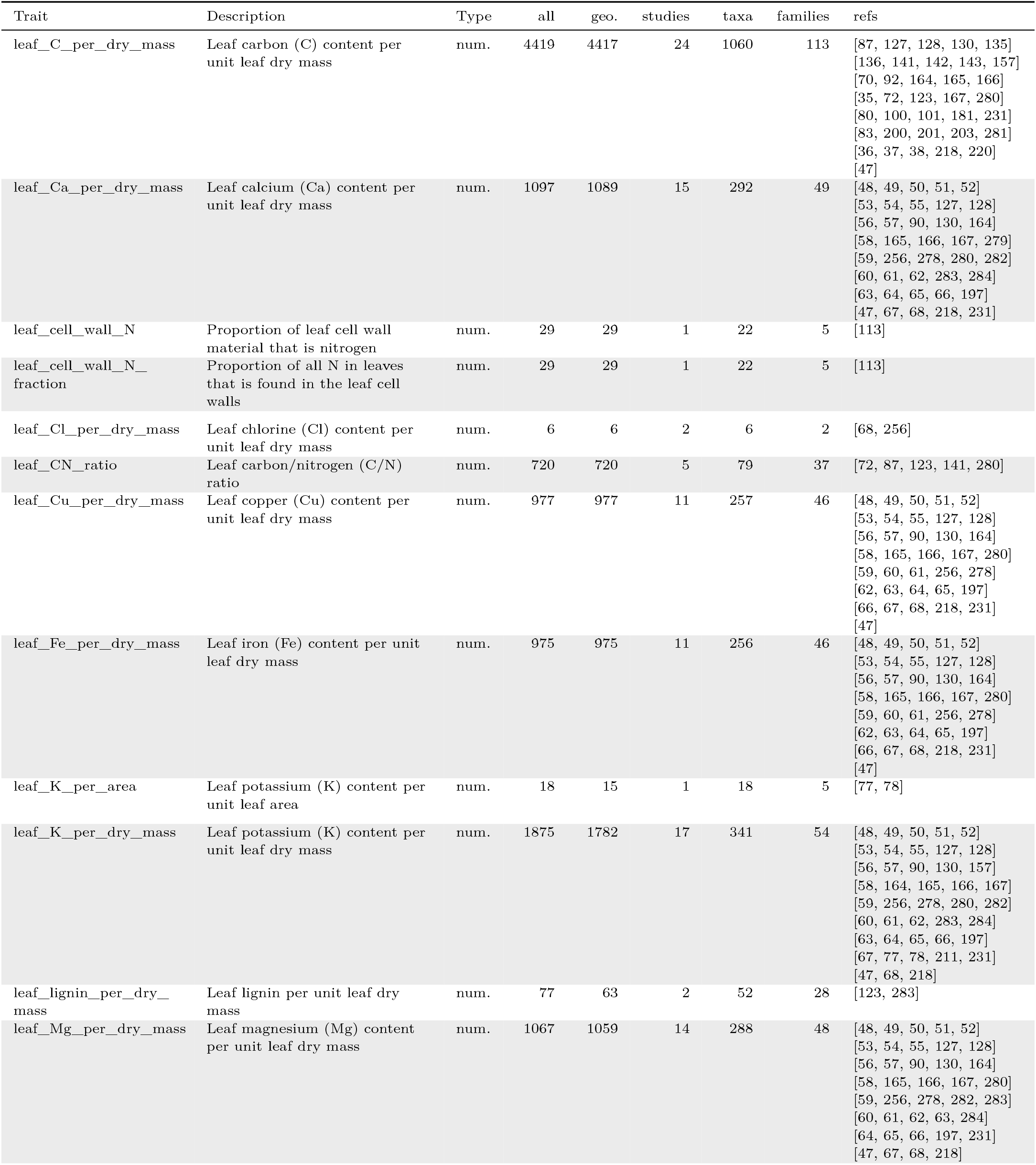
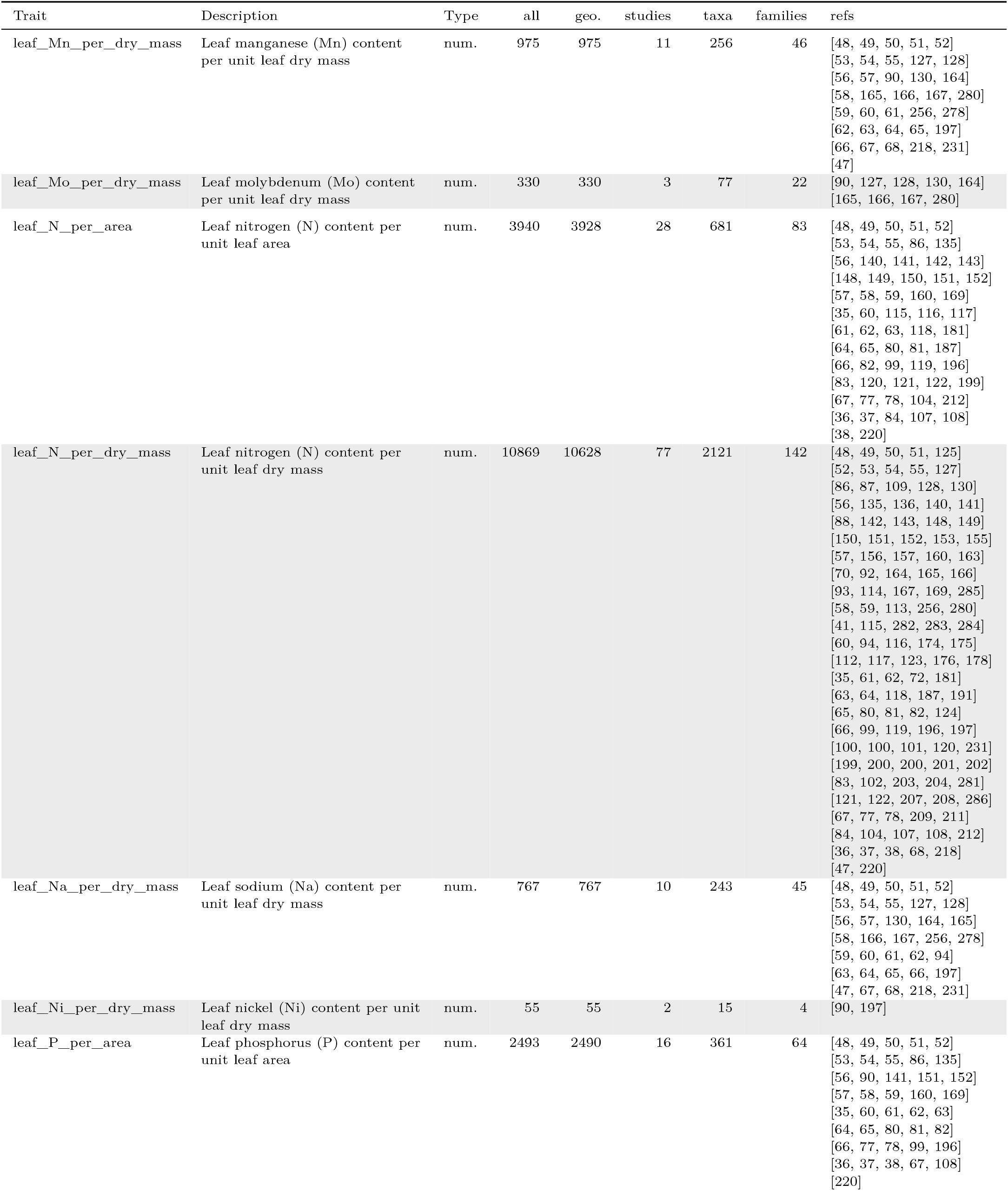
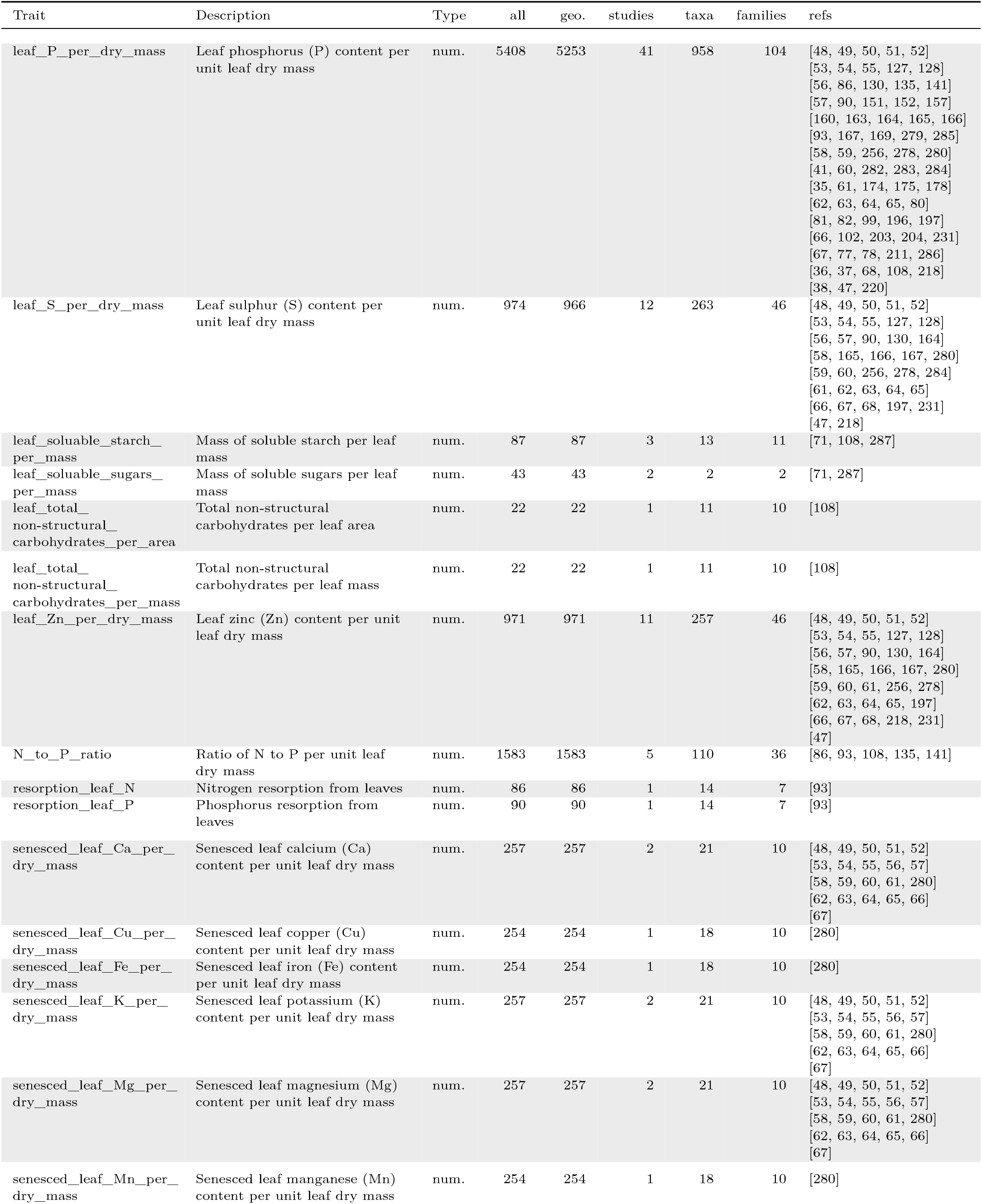
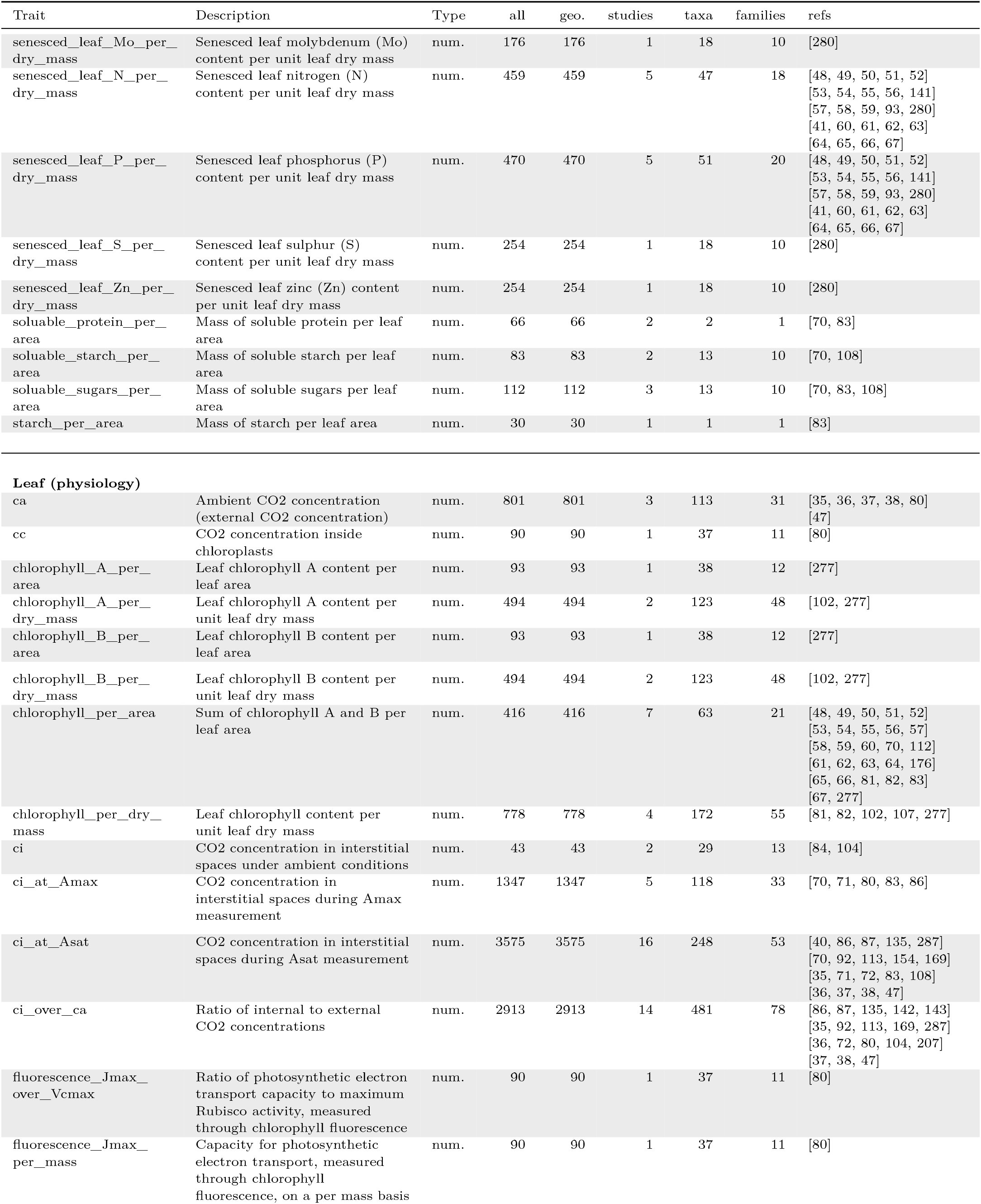
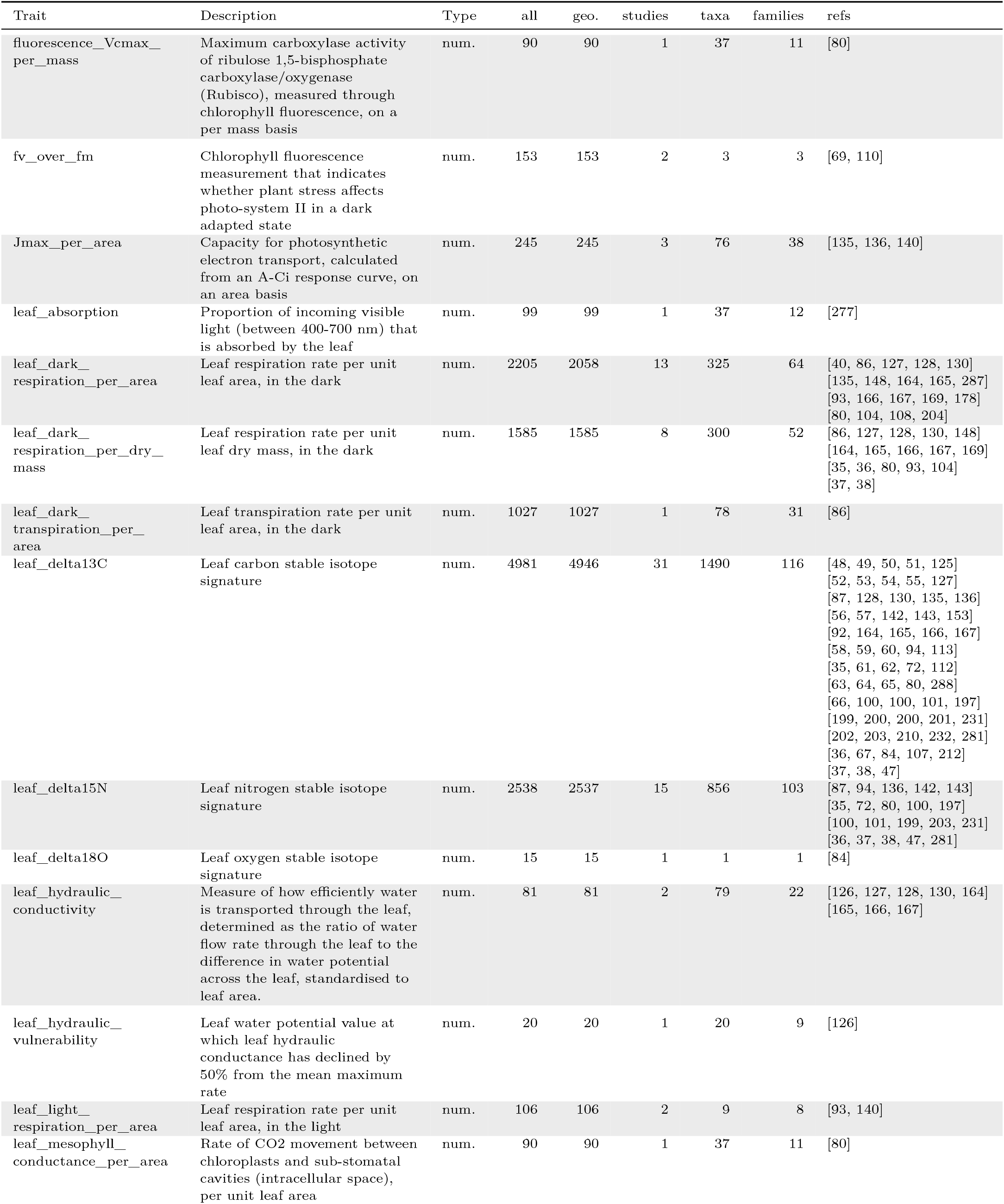
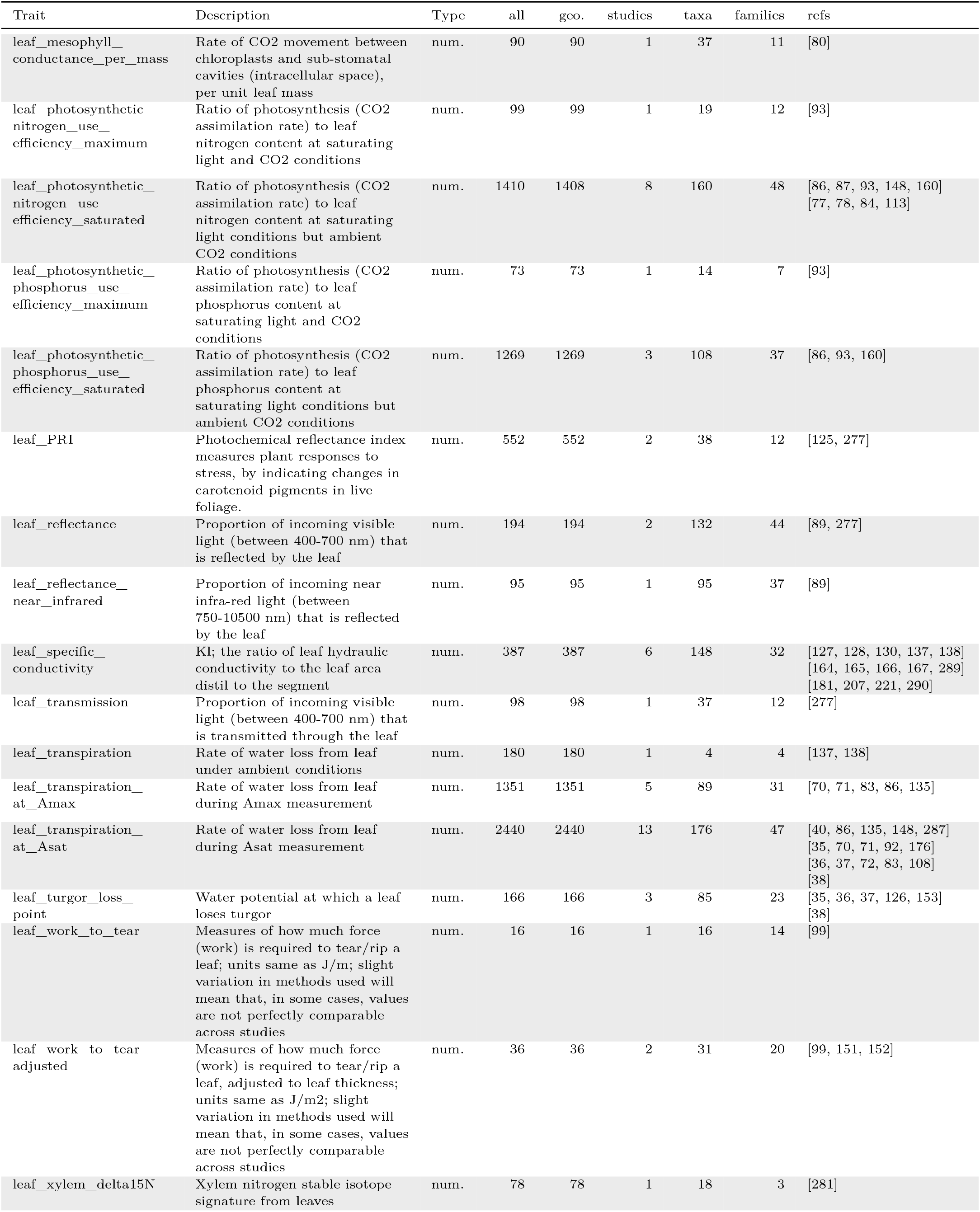
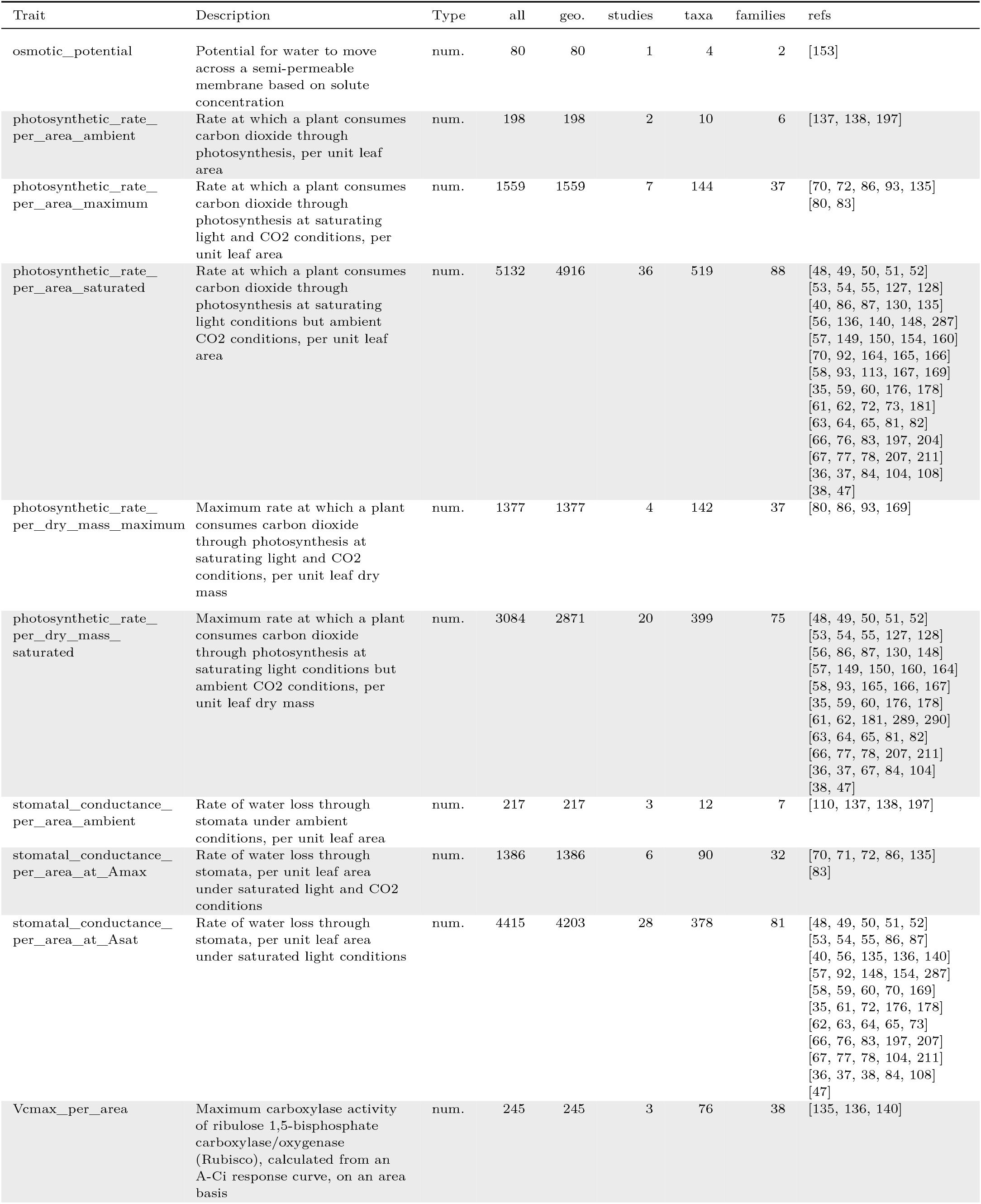
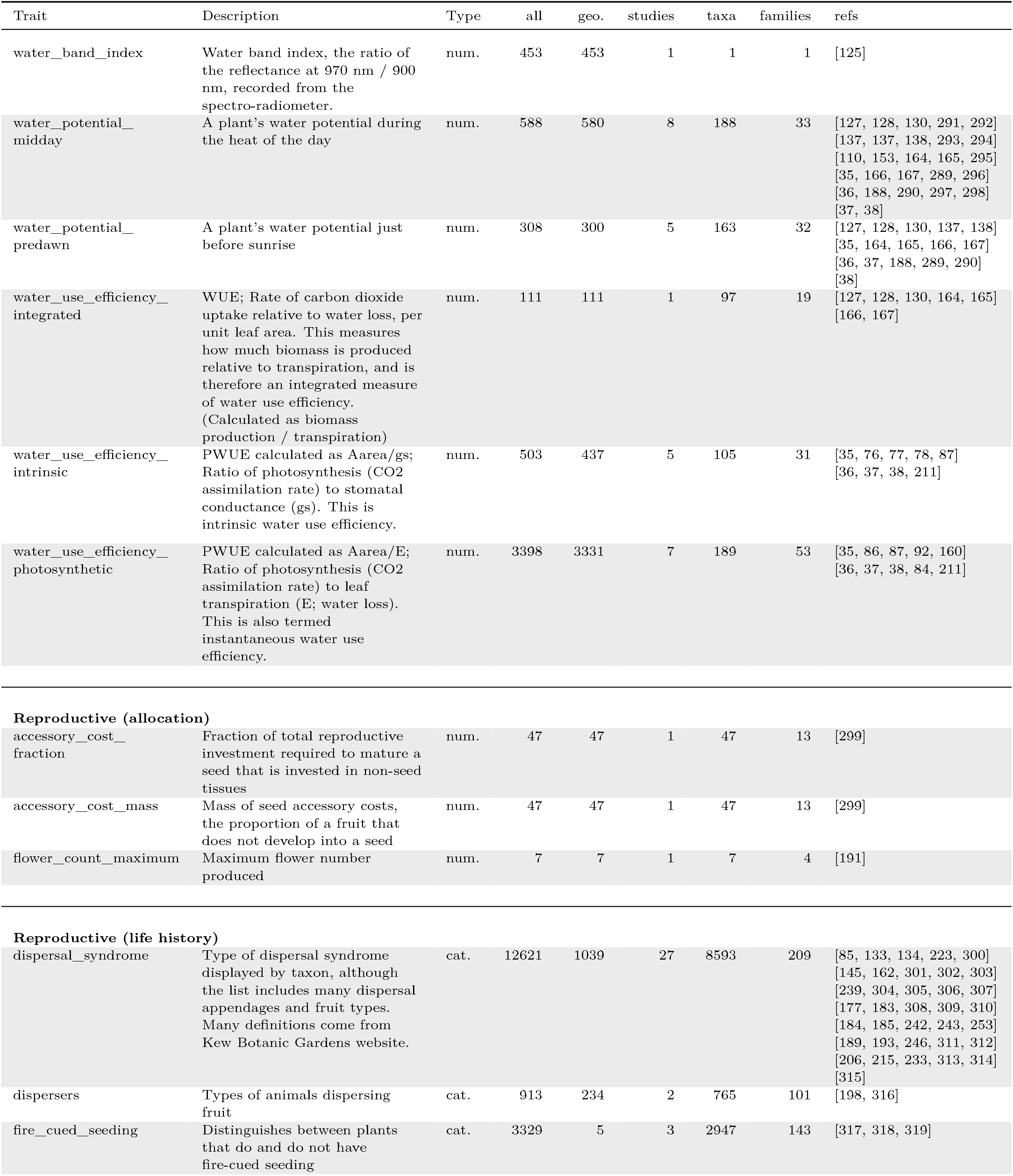
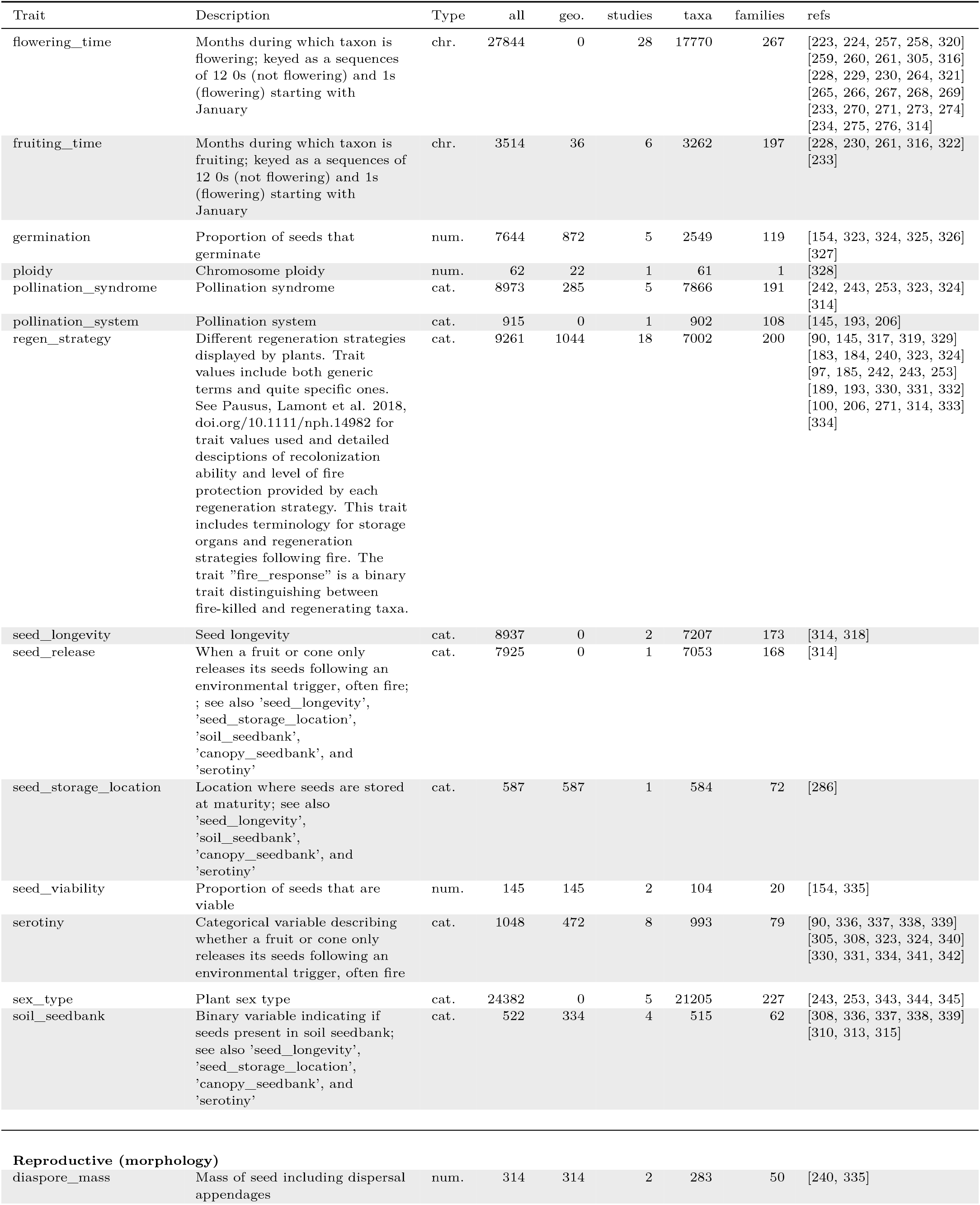
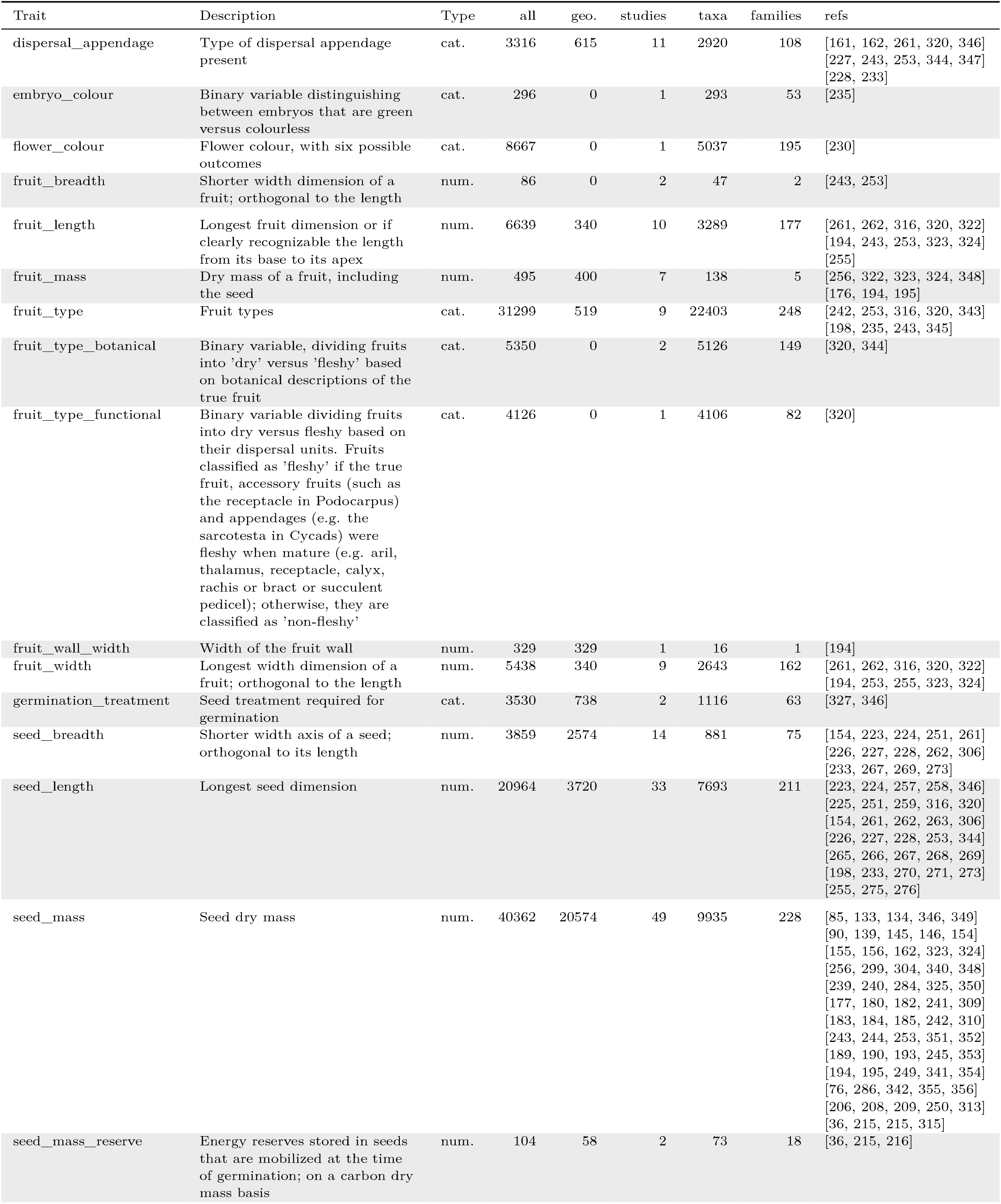
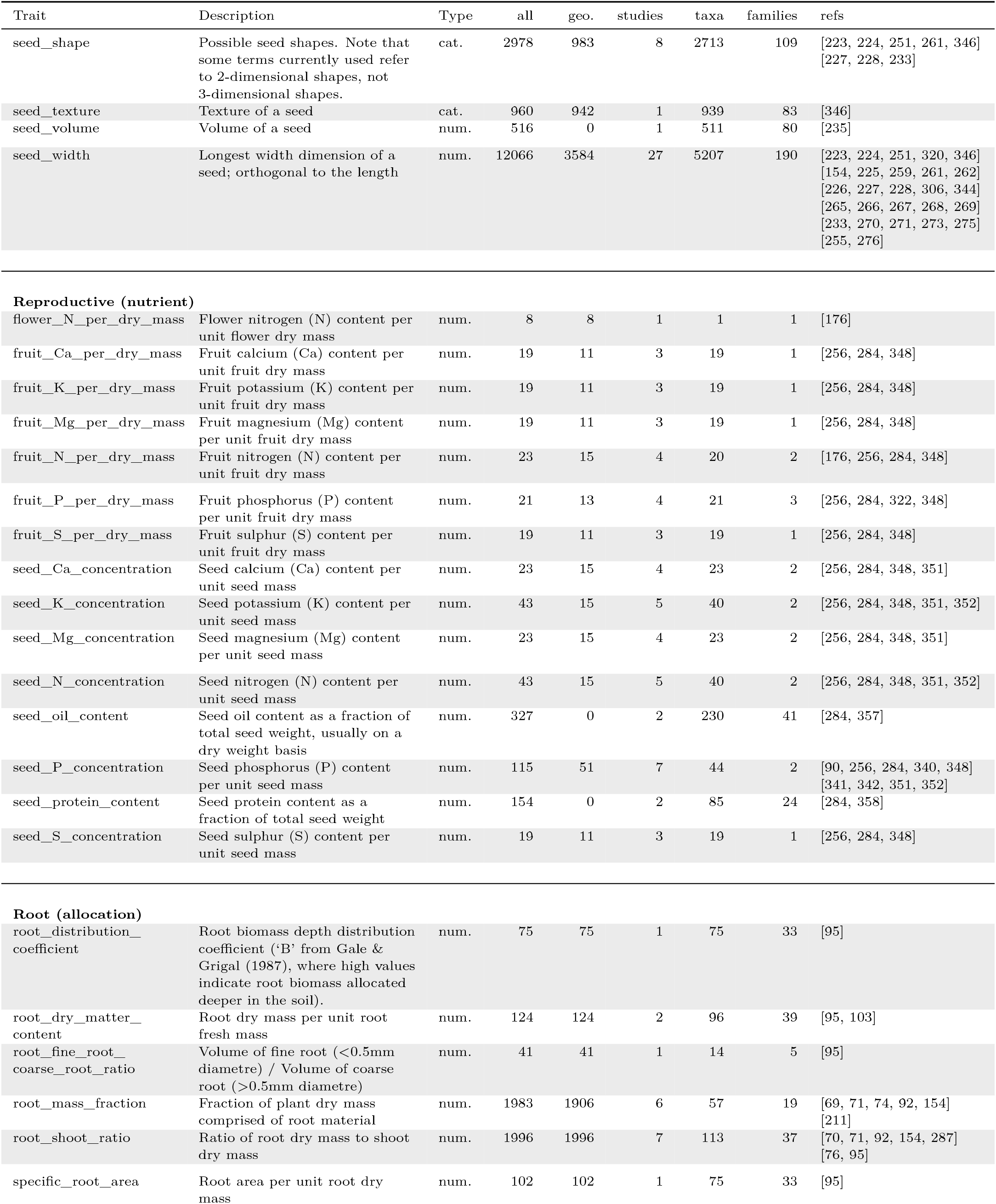
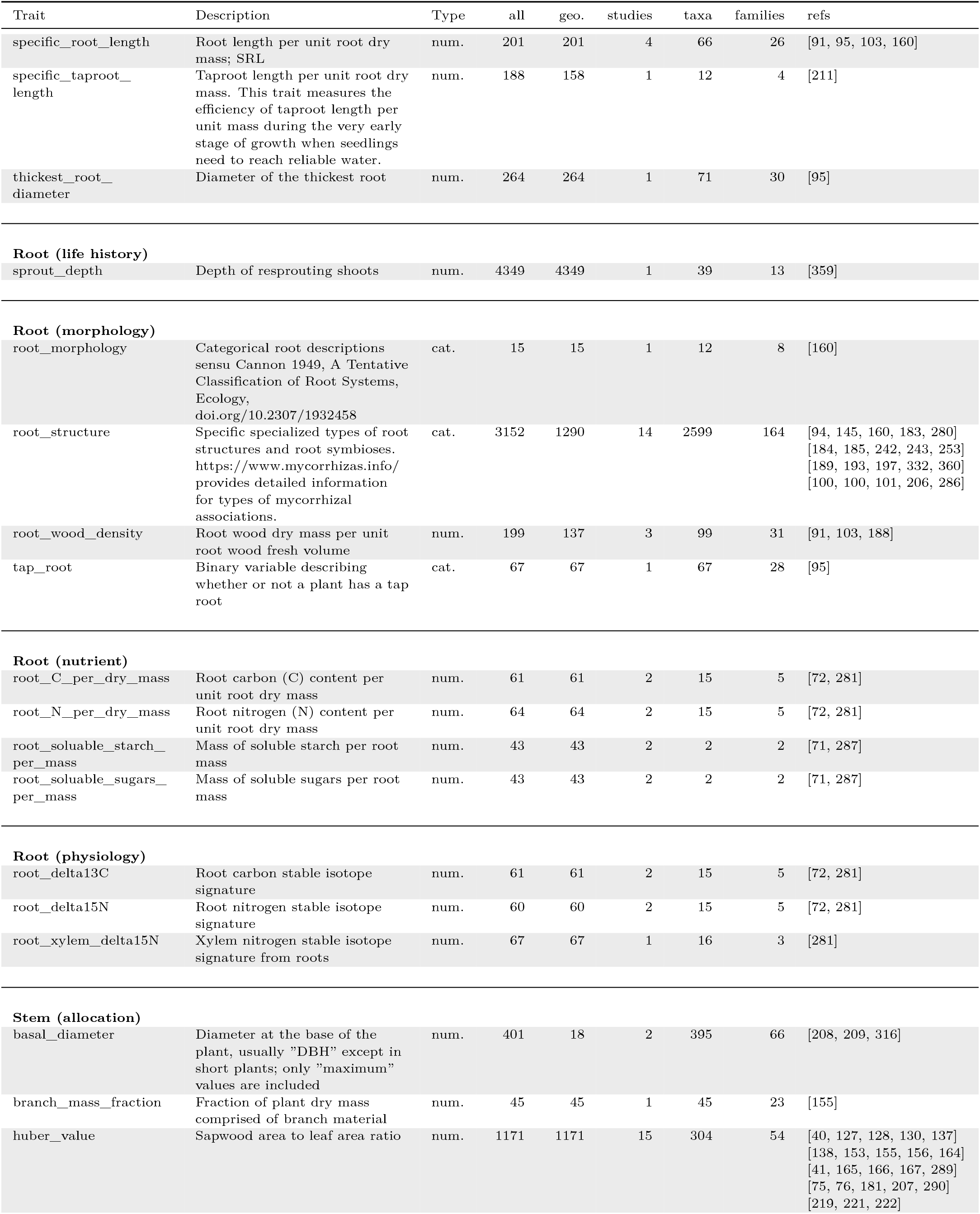
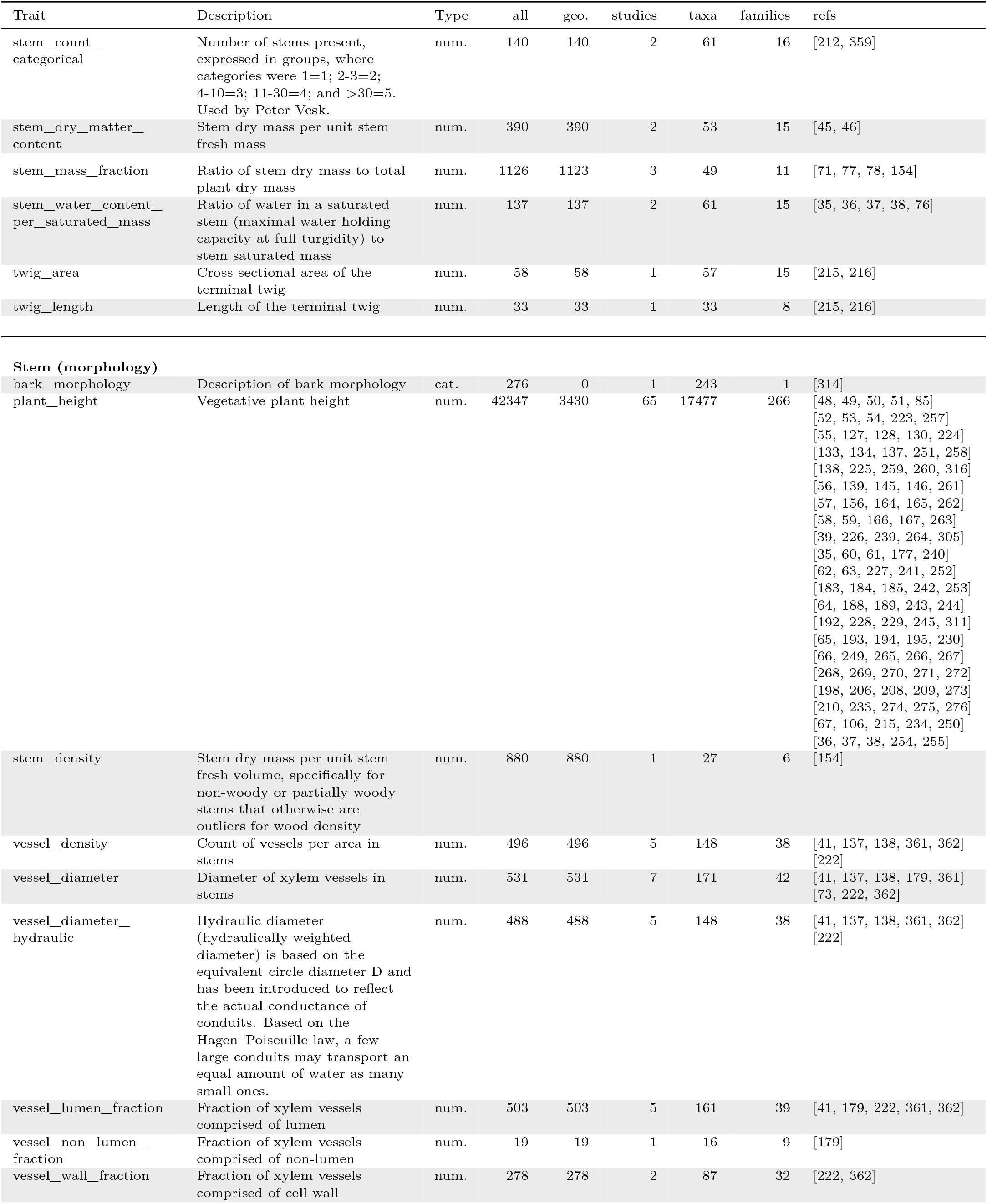
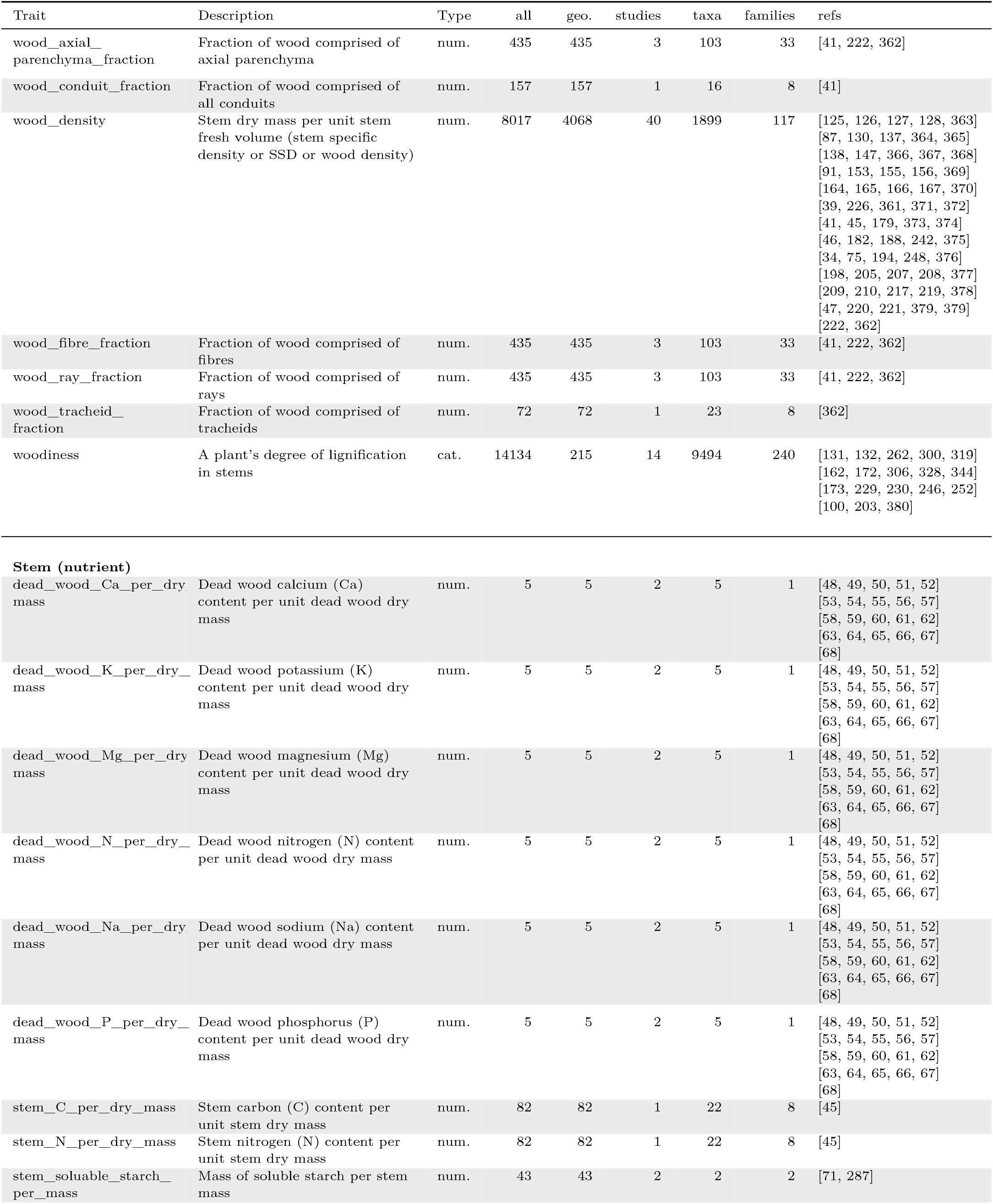
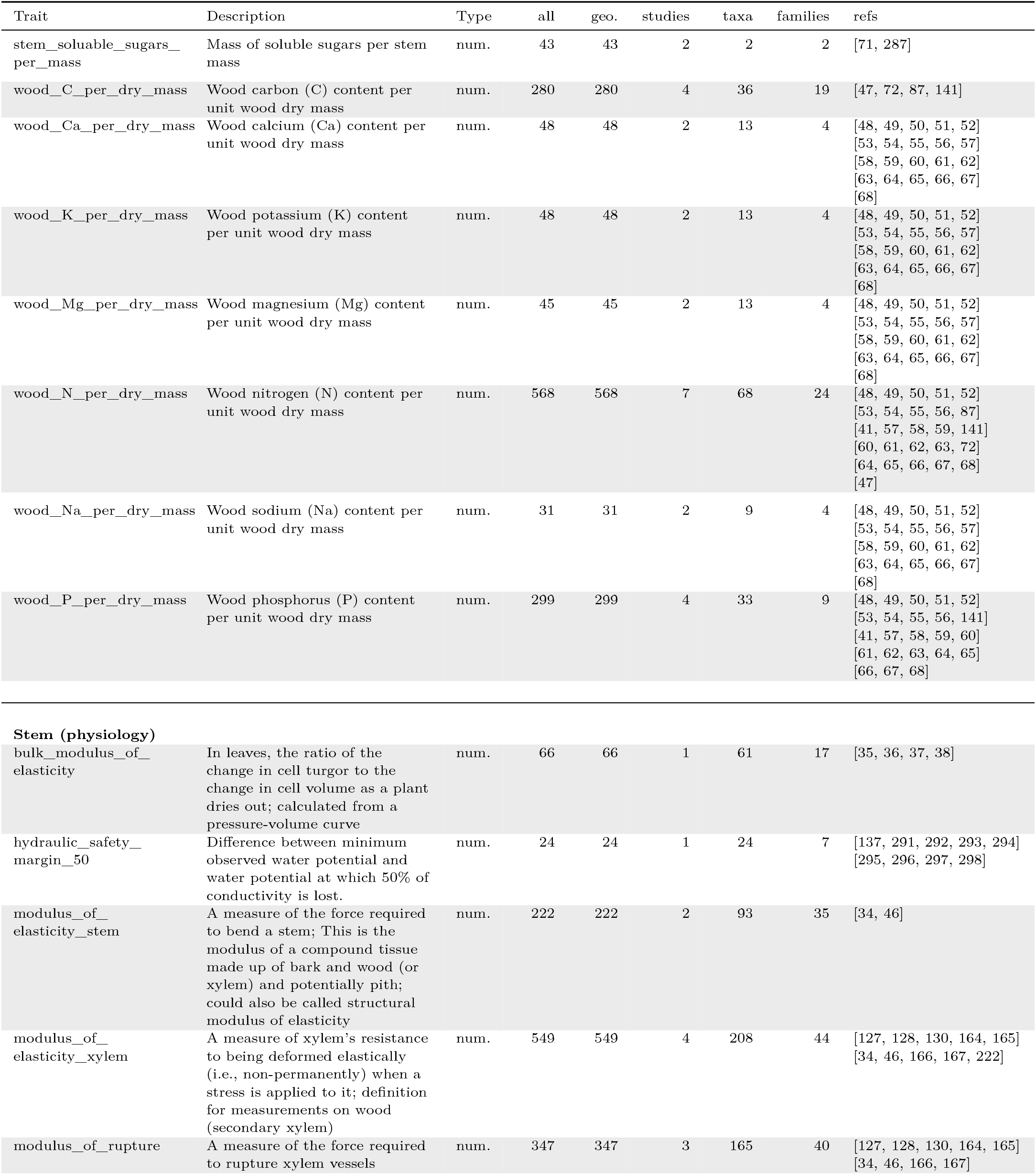
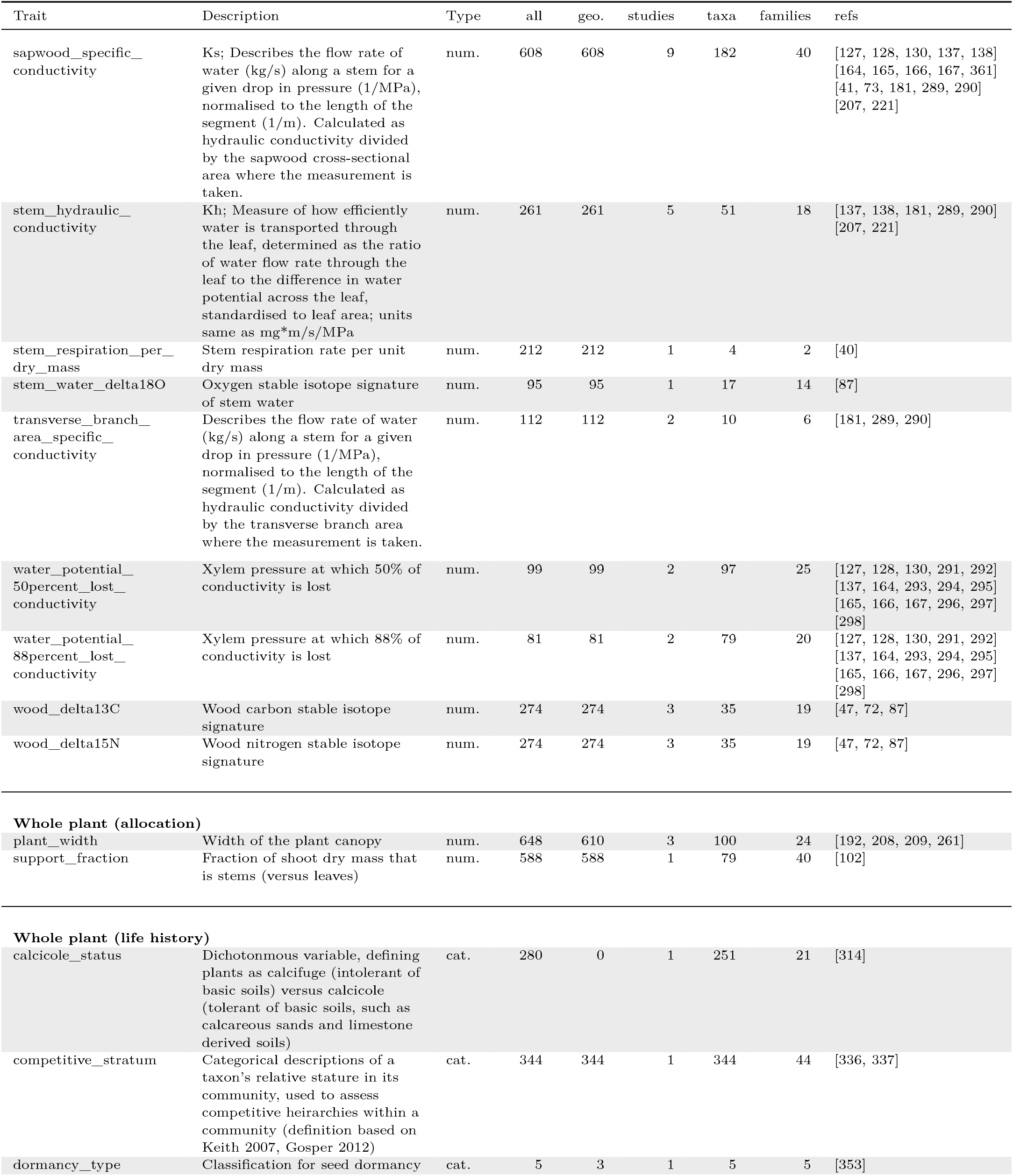
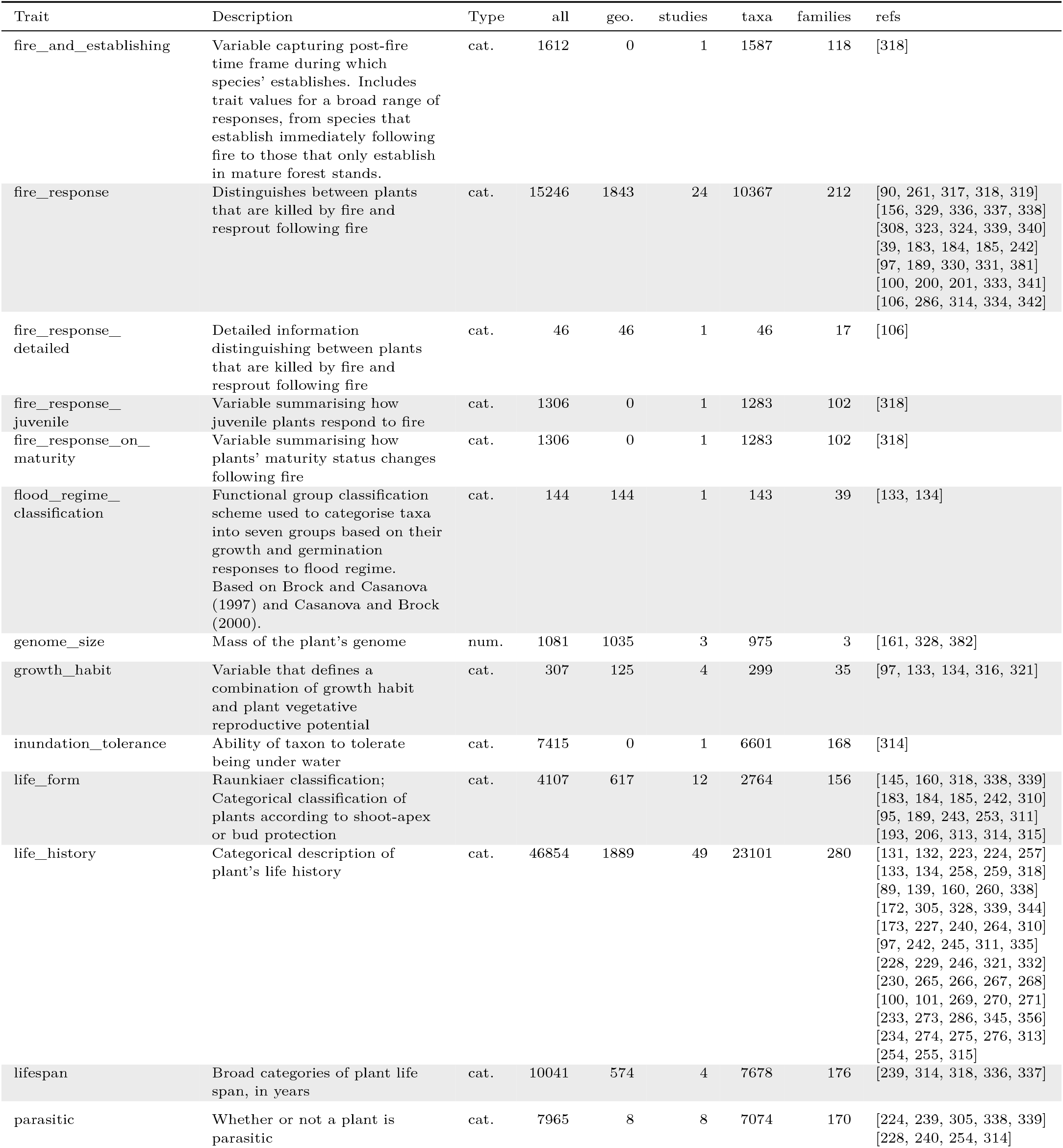
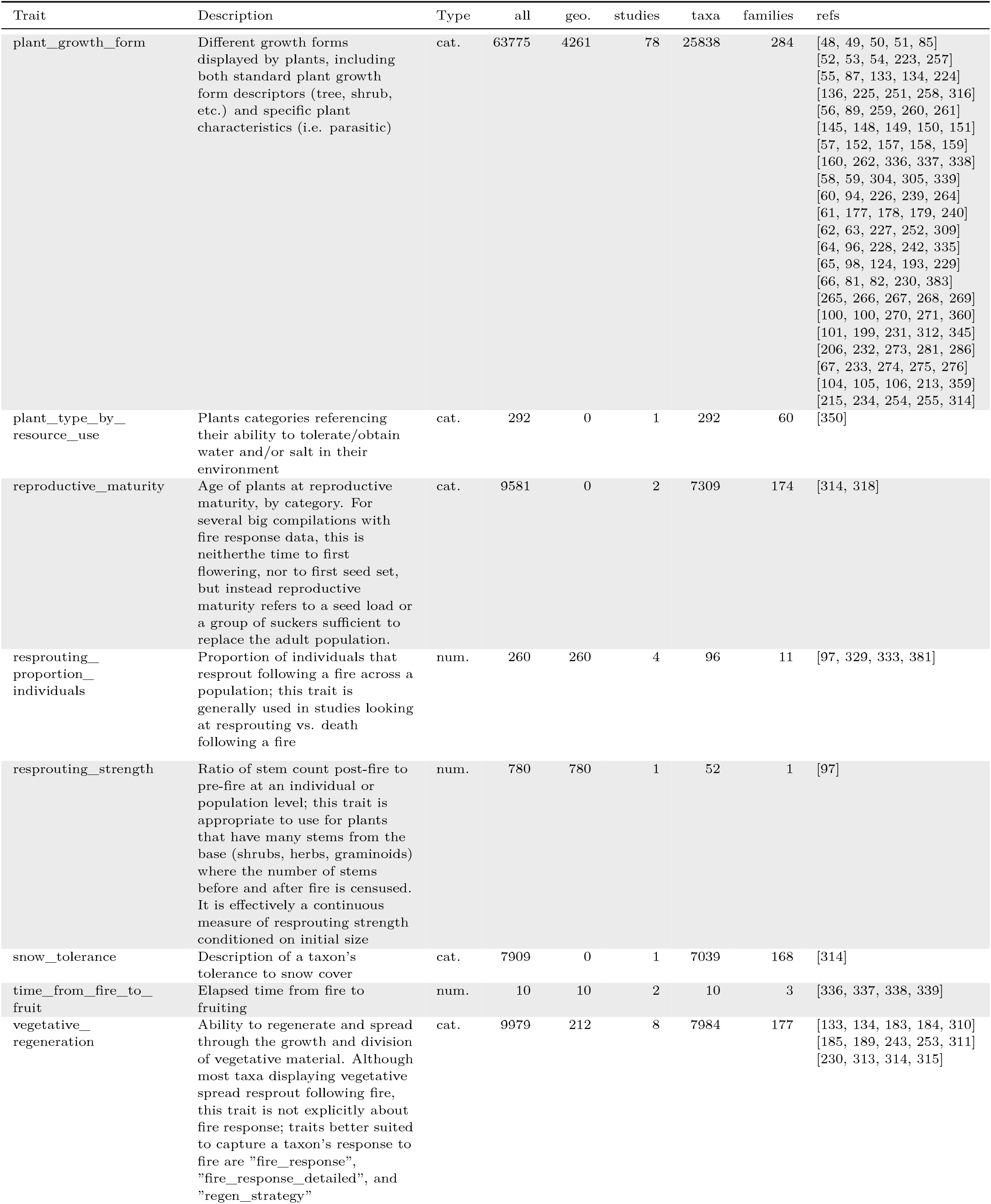
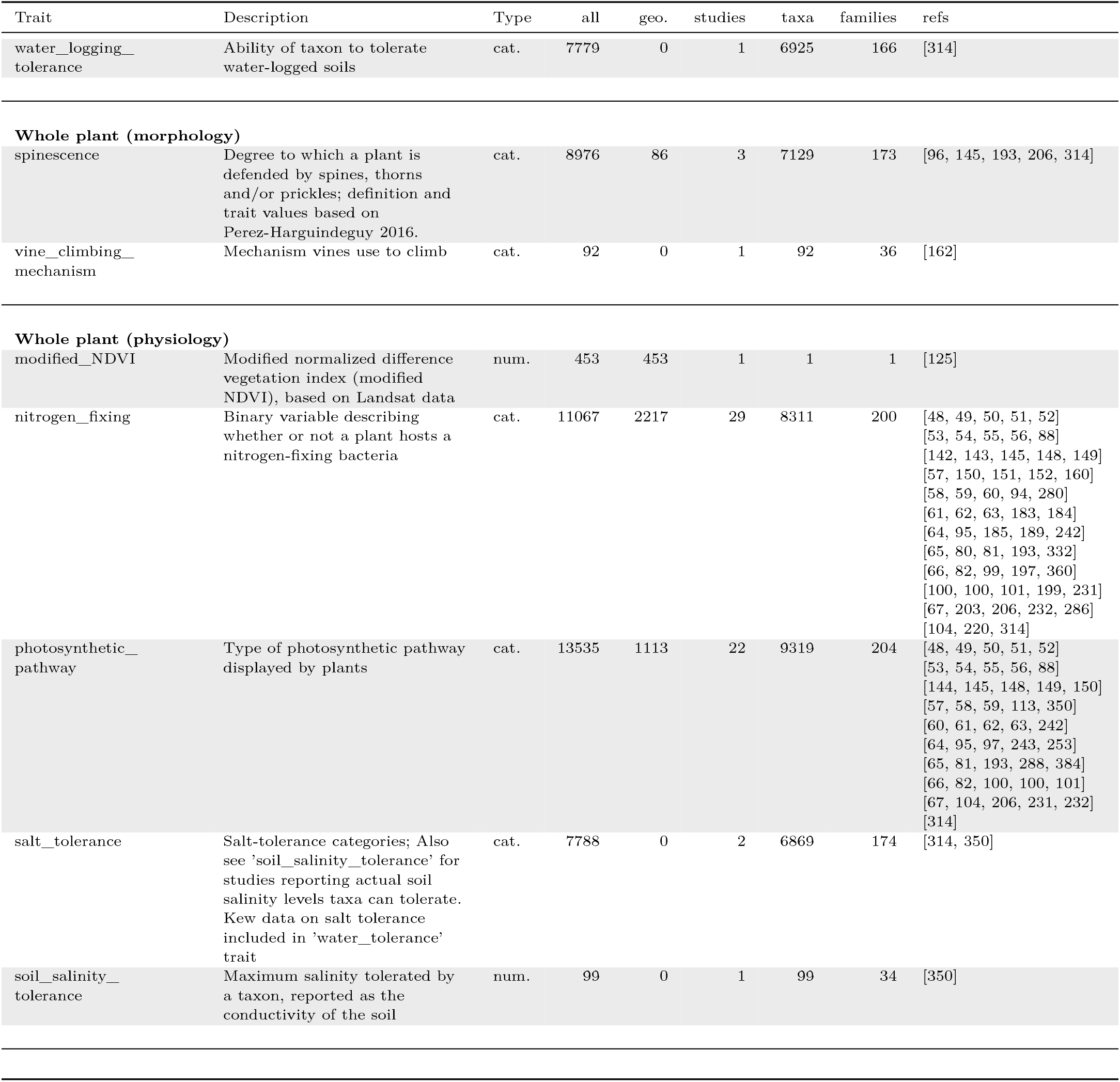
Details on all traits represented in version 2.1.0 of AusTraits. Note the count of studies is less than the number of references when studies are linked to multiple references.

There were substantial differences in coverage among different tissues and trait types, also with respect to number of geo-referenced points (Figure 2). The most common traits are non geo-referenced records from floras. Yet, geo-referenced records were available in several traits for more than 10% of the flora (Figure 2a).

**Figure 2:**
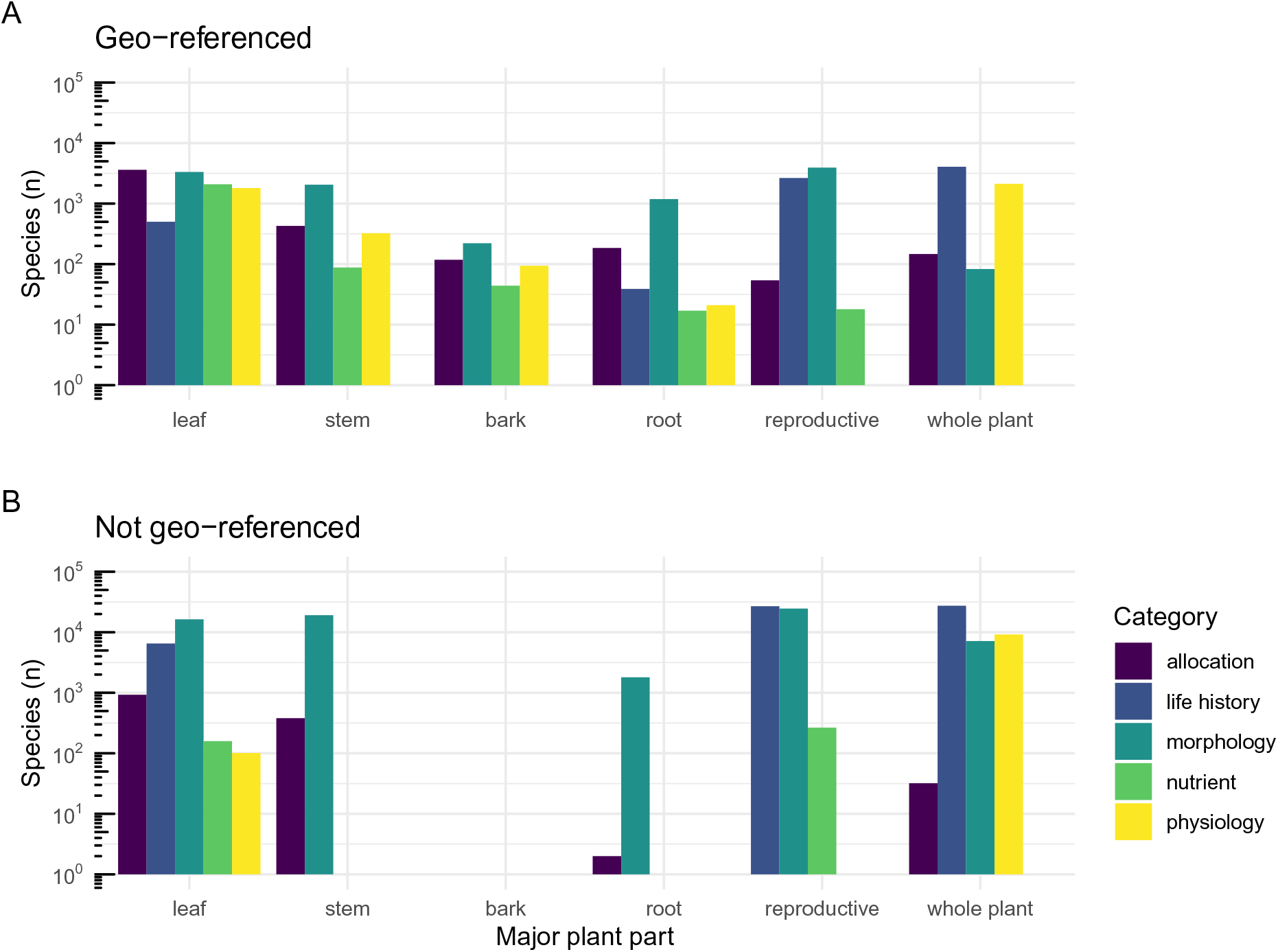
Number of taxa with trait records by plant tissue and trait category, for data that are (A) Geo-referenced, and (B) Not geo-referenced. Many records without a geo-reference come from botanical collections, such as floras.

We found that trait records were spread across the climate space of Australia (Figure 3a), as well as geographic locations (Figure 3b). As with most data, in Australia, the density of records was somewhat concentrated around cities or roads in remote regions, particularly for leaf traits.

**Figure 3:**
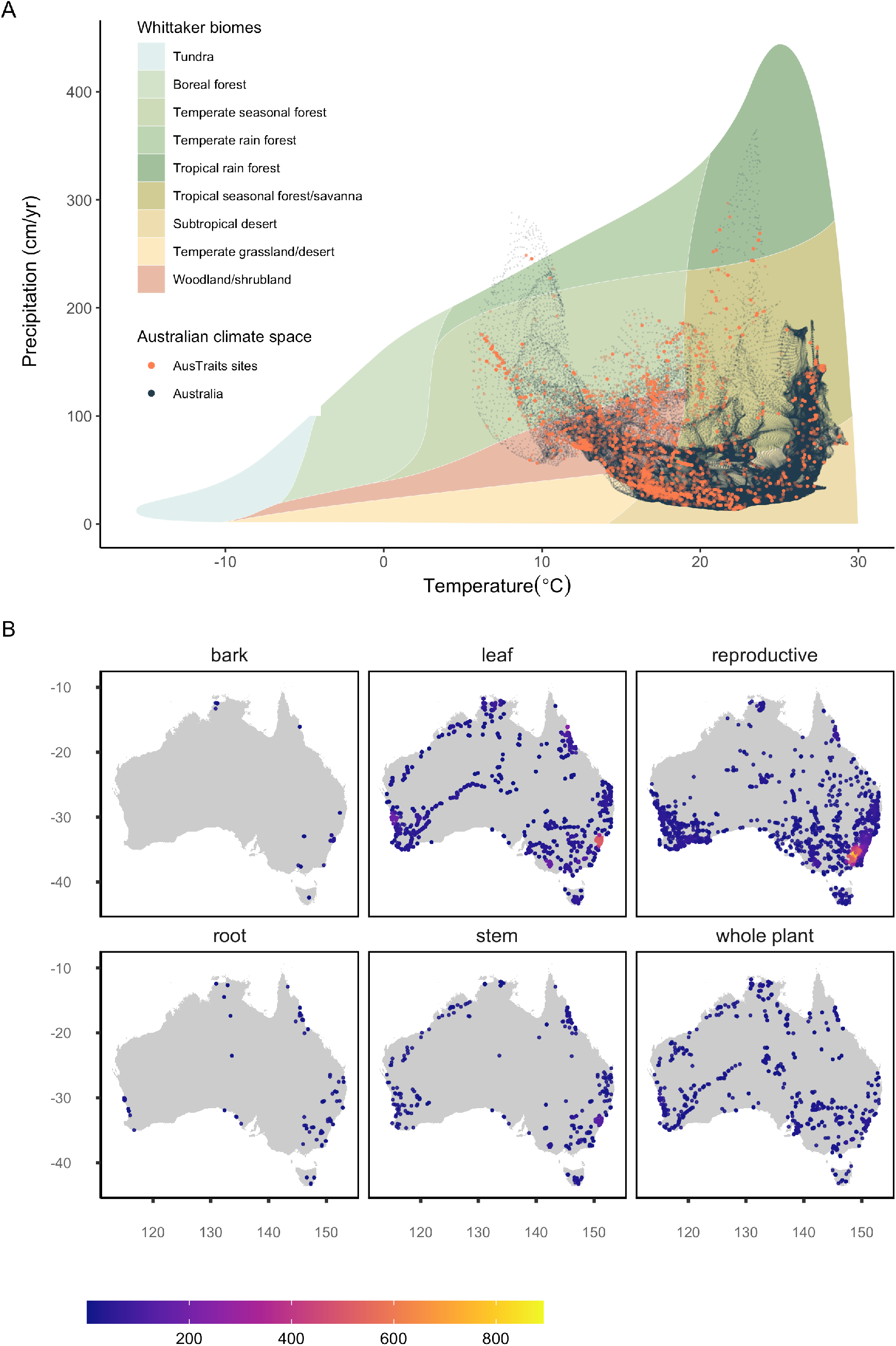
Coverage of geo-referenced trait records across Australian climatic and geographic space for traits in different categories. (A) AusTraits’ sites (orange) within Australia’s precipitation-temperature space (dark-grey) superimposed upon Whittaker’s classifctaion of majore biomes by climate [32]. Climate data were extracted at 10” resolution from WorldClim [33].(B) Locations of geo-referenced records for different plant tissues.

Figure 4 shows that overall coverage across a phylogenetic tree of Australian plant species is relatively unbiased, though there are some notable exceptions. One exception is for root traits, where taxa within Poaceae have large amounts of information available relative to other plant families. A cluster of taxa within the family Myrtaceae have little leaf information available, while reproductive information is limited for species near the base of the tree.

**Figure 4:**
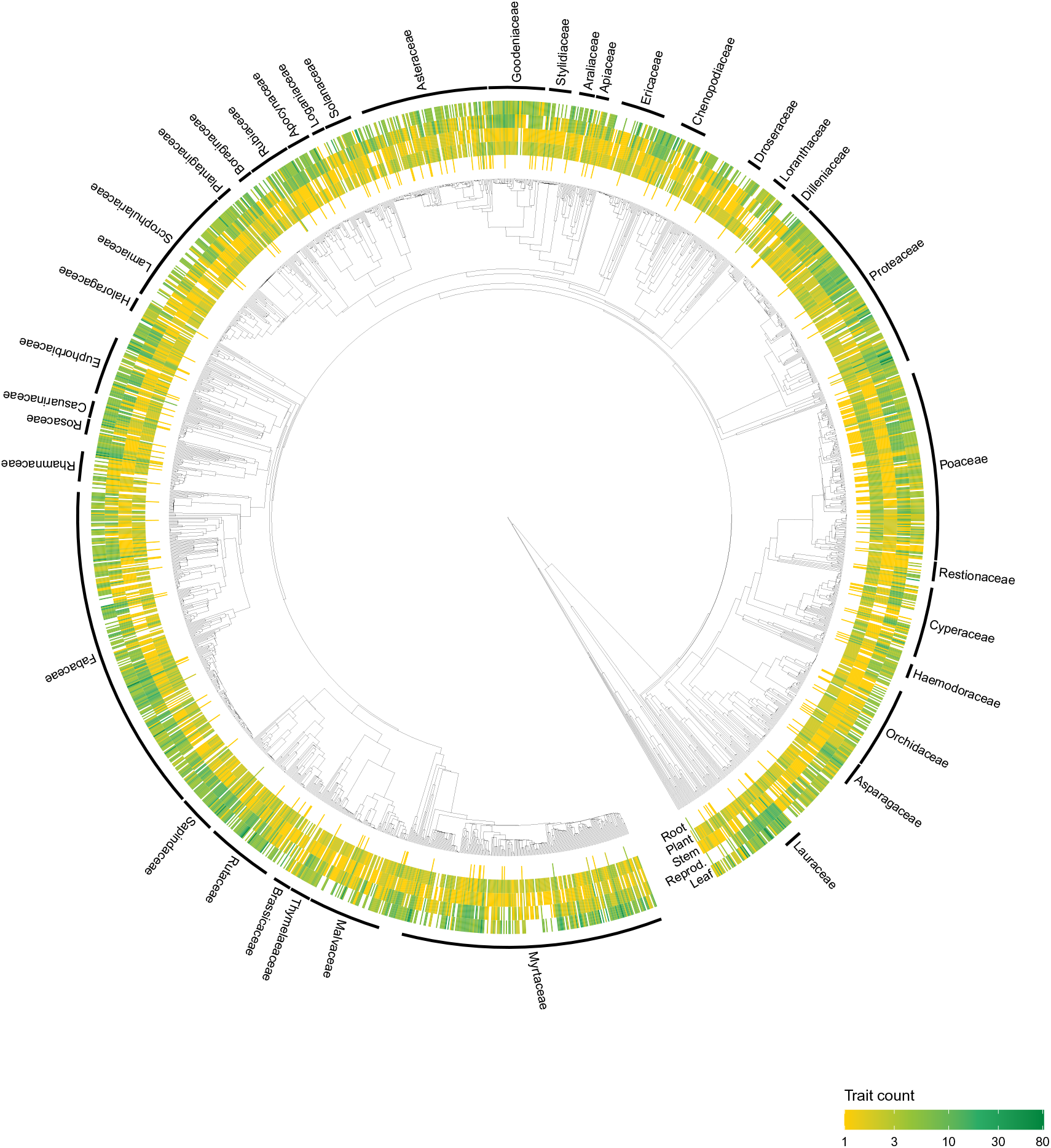
Phylogenetic distribution of trait data in AusTraits for a subset of 2000 randomly sampled taxa. The heatmap colour intensity denotes the number of traits measured within a family for each plant tissue. The most widespread family names (with more than ten taxa) are labelled on the edge of the tree.

Comparing coverage in AusTraits to the global database TRY, there were 72 traits overlapping. Of these, AusTraits tended to contain records for more taxa, but not always (Figure 5). Multiple traits had more than 10 times the number of taxa represented in AusTraits. However, there were more records in TRY for 22 traits, in particular physiological leaf traits. Many traits were not overlapping between the two databases (Figure 5). We noted that AusTraits includes more seed and fruit nutrient data; possibly reflecting the interest in Australia in understanding how fruit and seeds are provisioned in nutrient-depauperate environments. AusTraits includes more categorical values, especially variables documenting different components of species’ fire response strategies, reflecting the importance of fire in shaping Australian communities and the research to document different strategies species have evolved to succeed in fire-prone environments.

**Figure 5:**
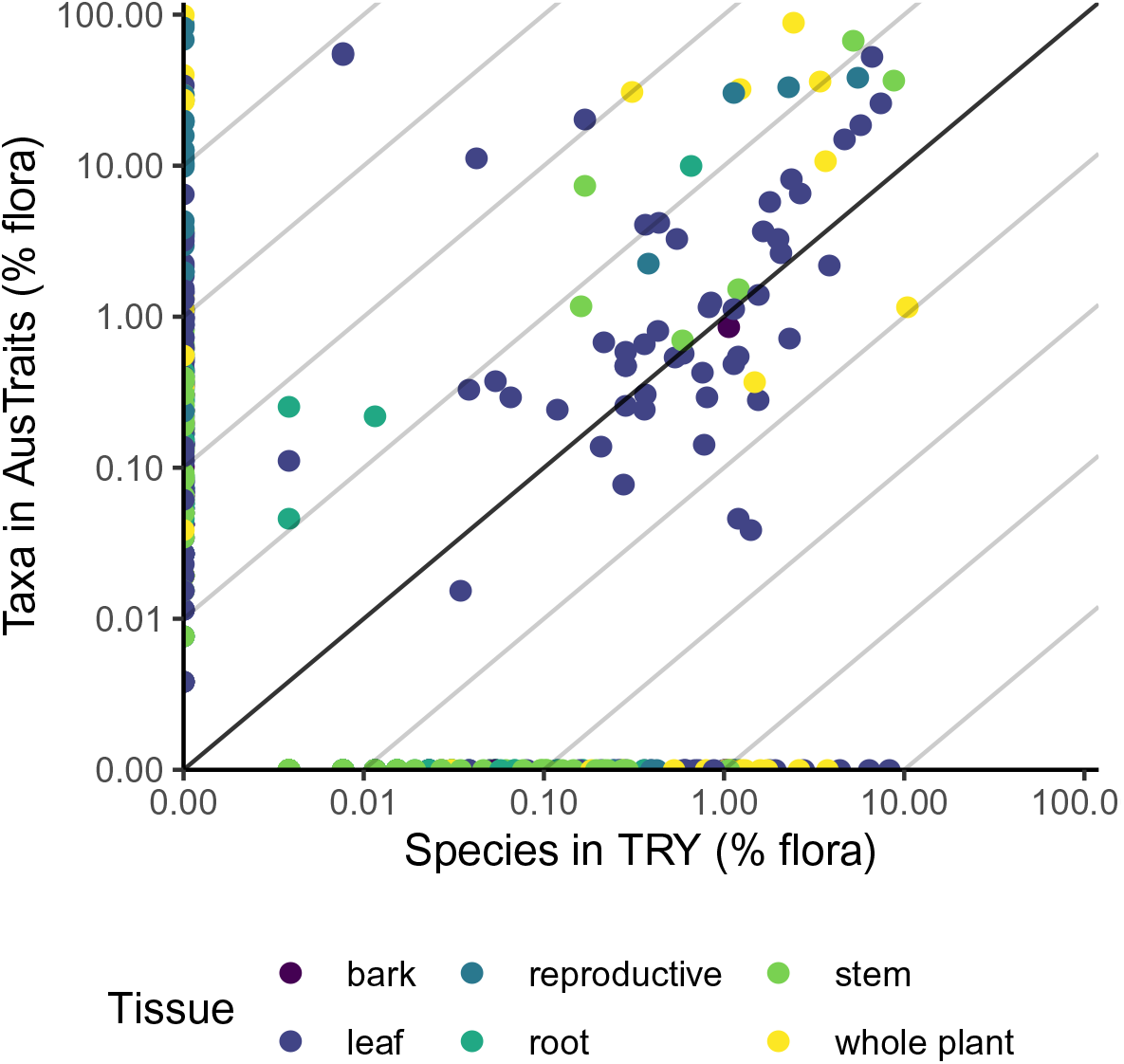
The number of taxa with trait records in AusTraits and global TRY database (accessed 28 May 2020). Each point shows a separate trait.

### Technical Validation

We implemented three strategies to maintain data quality. First, we conducted a detailed review of each source based on a bespoke report, showing all data and metadata, by both an AusTraits curator and the original contributor (where possible). Observations for each trait were plotted against all other values for the trait in AusTraits, allowing quick identification of outliers. Corrections suggested by contributors were combined back into AusTraits and made available with the next release.

Second, we implemented automated tests for each dataset, to confirm that values for continuous traits fall within the accepted range for the trait, and that values for categorical traits are on a list of accepted values maintained by the creators. Data that did not pass these tests were moved to a separate spreadsheet (“excluded_data”) that is also made available for use and review.

Third, we provide a pathway for user feedback. AusTraits is a community resource and we encourage engagement from users on maintaining the quality and usability of the dataset. As such, we welcome reporting of possible errors, as well as additions and edits to the online documentation for AusTraits that make using the existing data, or adding new data, easier for the community. Feedback can be posted as an issue directly at the project.

### Usage Notes

Each data release is available in multiple formats: first, as a compressed folder containing text files for each of the main components, second, as a compressed R object, enabling easy loading into R for those using that platform.

Using the taxon names aligned with the APC, data can be queried against location data from the Atlas of Living Australia. To create the phylogenetic tree in Figure 5, we pruned a master tree for all higher plants [28] using the package `V.PhyloMaker` [29] and visualising via `ggtree` [30]. To create Figure 3A, we used the package `plotbiomes` [31] to create the baseline plot of biomes.

### Code Availability

All code, raw and compiled data are hosted within GitHub repositories under the Trait Ecology and Evolution organisation (http://traitecoevo.github.io/austraits.build/). The archived material includes all data sources and code for rebuilding the compiled dataset. The code used to produce this paper is available at http://github.com/traitecoevo/austraits_ms. (All code will be made available prior to final publication.)

## Acknowledgements

This work was supported by fellowship grants from Australian Research Council to Falster (FT160100113), Gallagher (DE170100208) and Wright (FT100100910) and funding from Macquarie University to Gallagher, and the Australian Research Data Commons via their “Transformation data collections” program. We gratefully acknowledge input from the following persons who contributed to data collection Anna Monro, Sophia Amini, Julian Ash, Tara Boreham, Willi A. Brand, Amber Briggs, John Brock, Don Bulter, Robert Chinnock, Peter Clarke, Derek Clayton, Steven Clemants, Harold Trevor Clifford, Michelle Cochrane, Bronwyn Collins, Alessandro Conti, Wendy Cooper, William Cooper, Ian Cowie, Lyn Craven, Ian Davidson, Derek Eamus, Judy Egan, Chris Fahey, Paul Irwin Forster, John Foster, Tony French, Allison Frith, Ronald Gardiner, Ethel Goble-Garratt, Peter Grubb, Chris Guinane, TJ Hall, Monique Hallet, Tammy Haslehurst, Foteini Hassiotou, John Herbohn, Peter Hocking, Jing Hu, Kate Hughes, Muhammad Islam, Ian Kealley, Greg Keighery, James Kirkpatrick, Kirsten Knox, Luka Kovac, Kaely Kreger, John Kuo, Martin Lambert, Dana Lanceman, Michael Lawes, Claire Laws, Emma Laxton, Liz Lindsay, Daniel Montoya Londono, Christiane Ludwig, Ian Lunt, Mary Maconochie, Karen Marais, Bruce Maslin, Riah Mason, Richard Mazanec, Kate McClenahan, Elissa McFarlane, Huw Morgan, Peter Myerscough, Des Nelson, Dominic Neyland, Mike Olsen, Jacob McC. Overton, Paula Peeters, George Perry, Aaron Phillips, Loren Pollitt, Rob Polmear, Aina Price, Thomas Pyne, R.J.Williams, Barbara Rice, Jessica L. Rigg, Bryan Roberts, Miguel de Salas, Anna Salomaa, Inge Schulze, Waltraud Schulze, Andrew John Scott, Alison Shapcott, Luke Shoo, Anne Sjostrom, Santiago Soliveres, Amanda Spooner, George Stewart, Jan Suda, Catherine Tait, Daniel Taylor, Ian Thompson, Hellmut R. Toelken, Malcolm Trudgen, W.E Westman, Erica Williams, Kathryn Willis, J. Bastow Wilson, Jian Yen. We acknowledge the work of all Australian taxonomists and their supporting institutions, whose long-term work on describing the flora has provided a rich source of data for AusTraits.

## Author contributions

RVG, IJW conceived the original idea; RVG, EHW, CB, SA collated data from primary sources; DSF developed the workflow for the harmonising of data and led all coding; EHW, DI, SCA, JL contributed to coding; EHW, SCA, CB, JL error-checked trait observations; DI developed figures for the paper; DSF, RVG, DI, EHW wrote the first draft of the paper. All other authors contributed the raw data and metadata underpinning the resource, reviewed the harmonised data for errors, and reviewed the final paper for publication.

## Overview

AusTraits harmonises data on 375 traits from 264 different sources, including field campaigns, published literature, taxonomic monographs, and individual taxon descriptions.

This document provides information on the structure of AusTraits and corresponds to version 2.1.0 of the dataset.

